# Preconception e-nicotine impairs airway development and progenitor proliferation across generations in Drosophila *Melanogaster*

**DOI:** 10.64898/2026.06.08.730871

**Authors:** Leizhi Shi, Qiaoyan Tan, Fan Xu, Hanna Angstmann, Judith Bossen, Sandra Uzenabor, Sven Künzel, Daniel Meissner, Birte Kemper-Ehrhardt, Göksel Dogan, Beate Hoeschler, Sai Sneha Priya Nemani, Klaus F. Rabe, Cecilie Svanes, Thomas Roeder, Susanne Krauss-Etschmann

## Abstract

Electronic nicotine delivery devices are increasingly used worldwide, raising concern about the potential effects of nicotine exposure during fetal and early postnatal development. However, evidence regarding the effects of vaping during pregnancy remains limited, and the consequences of exposure before conception are largely unknown. Here, we investigated whether maternal nicotine vaping prior to conception alters airway development using the fruit fly *Drosophila melanogaster*. Offspring of nicotine-exposed mothers exhibited abnormal airway architecture, epithelial remodelling, and impaired terminal branching. These defects were accompanied by reduced proliferation of airway progenitor cells without increased apoptosis, resulting in decreased hypoxia tolerance, altered activity patterns, and reduced lifespan. Transcriptomic and functional analyses revealed suppression of innate immune signalling mediated by nuclear factor kappa-B (NF-κB). Notably, knockdown of its regulator *Relish* reproduced key phenotypes. Developmental abnormalities persisted across generations, with their severity progressively declining. Together, these findings identify maternal preconception nicotine exposure as a previously unrecognized determinant of airway development and provide insight into early origins of chronic airway disease.

**Teaser:** Preconception e-nicotine exposure shapes airway development, structural and physiological abnormalities across generations in Drosophila.

## Introduction

The use of electronic cigarettes (e-cigarettes) has expanded rapidly from North America to Europe and Asia, particularly among adolescents and young adults, including women of reproductive age (*1, 2*). This trend has raised concern about the health consequences of nicotine exposure during critical developmental periods. Early life environmental exposures are known to influence later physiological function and disease susceptibility (*3–5*), making it important to determine whether nicotine exposure during sensitive developmental windows affects offspring respiratory health.

Maternal e-cigarette use during pregnancy is increasingly prevalent, with global estimates ranging from 0.5 % to 15 % (*6*). However, epidemiological evidence linking maternal e-cigarette exposure to respiratory outcomes in offspring remains limited. Nicotine, a key component of e-cigarettes, can reach concentrations of up to 6 % (*7*), often exceeding those of combustion cigarettes (*8*), and accumulates in fetal tissues at biologically relevant levels (*9*). Human studies examining offspring outcomes are scarce, although several reports have described associations with lower birth weight, preterm birth, and small-for-gestational-age infants (*10–13*). Interpretation remains challenging because concurrent use of combustible cigarettes may confound observed associations (*14*). Notably, vulnerability extends beyond *in utero* exposure. Emerging evidence suggests that parental puberty and the preconception period represent additional critical windows during which smoking can alter germline epigenetic programming and shape offspring respiratory trajectories (*15*). A substantial body of epidemiological and experimental evidence demonstrates that prenatal exposure to nicotine or tobacco smoke disrupts airway development and increases susceptibility to respiratory dysfunction in offspring (*16–18*). Structural abnormalities, including airway wall thickening, impaired epithelial organization, and altered branching morphogenesis, have been consistently reported in both human and animal studies following maternal smoking during pregnancy (*18–21*). These findings have raised increasing concerns about the potential long-term and intergenerational health consequences of nicotine exposure during early developmental windows, including fetal and early postnatal development (*22–24*). Whether exposure occurring prior to conception—particularly during adolescence or early adulthood—can similarly compromise airway development in the next generation remains largely unknown. Lung development is a highly dynamic process characterized by rapid cellular proliferation, lineage differentiation, and coordinated branching morphogenesis and alveolarization. Experimental studies in mammalian models have shown that maternal e-cigarette exposure during pregnancy induces emphysema-like lung injury in offspring and disrupts developmental pathways central to lung development (*25–29*). Importantly, whether adolescent vaping prior to conception also compromises airway epithelial organization, terminal branching, and progenitor cell pools in offspring remains unresolved and represents a critical knowledge gap. Addressing this question is essential to define the inter- and transgenerational impact of maternal vaping on offspring airway health.

The fruit fly *Drosophila melanogaster* provides a genetically tractable airway system that recapitulates conserved principles of branching morphogenesis, airway epithelial organization, and alveolar-like cell specialization (*30*). It therefore represents a robust platform to interrogate the consequences of developmental perturbations across generations. Foundational studies have established quantitative frameworks for tracheal morphometric analyses (*31*) and delineated key cellular and molecular programs governing progenitor cell activation and airway development (*32*). In addition, the short generation time of the fruit fly together with precisely controllable exposures (*33*) to aerosols, enables systematic dissection of inter- and transgenerational effects that are difficult to study in humans due to ethical and temporal constraints. Here, we leverage this model to investigate whether maternal e-nicotine exposure prior to conception disrupts airway development and induces heritable alterations in offspring respiratory structure and function, and to define the underlying cellular and molecular mechanisms in a sex-specific manner.

## Results

### Preconception nicotine vaping alters offspring airway morphology and causes epithelial remodelling

To determine whether maternal e-nicotine exposure prior to conception affects airway development in the next generation, we first assessed its effects on physiological consequences. model. Exposure to vaporized nicotine, increased mortality and reduced lifespan in both female and male parental flies (Supplementary Fig. 1 and Supplementary Fig. 2A–B). Next, we examined the morphology of the dorsal trunk in the tracheal segments (T8–T9) of F_1_ larvae. Representative images revealed structural abnormalities in offspring from nicotine-exposed mothers, most notably focal narrowing of the dorsal trunk (Fig. 1A–D; red arrows). These abnormalities were observed at a markedly higher frequency in the nicotine-exposed group than in controls (Fig. 1E). Furthermore, the length of the dorsal trunk in the ninth tracheal segment (T9) was significantly reduced in female offspring from e-nicotine exposed mothers, whereas no significant difference was detected in males (Fig. 1F). In addition, secondary branch length was significantly reduced in the e-nicotine exposure group (Fig. 1G), while the number of secondary branches remained unchanged (Fig. 1H), suggesting that nicotine exposure primarily affects branch outgrowth rather than branch initiation.

**Figure 1:**
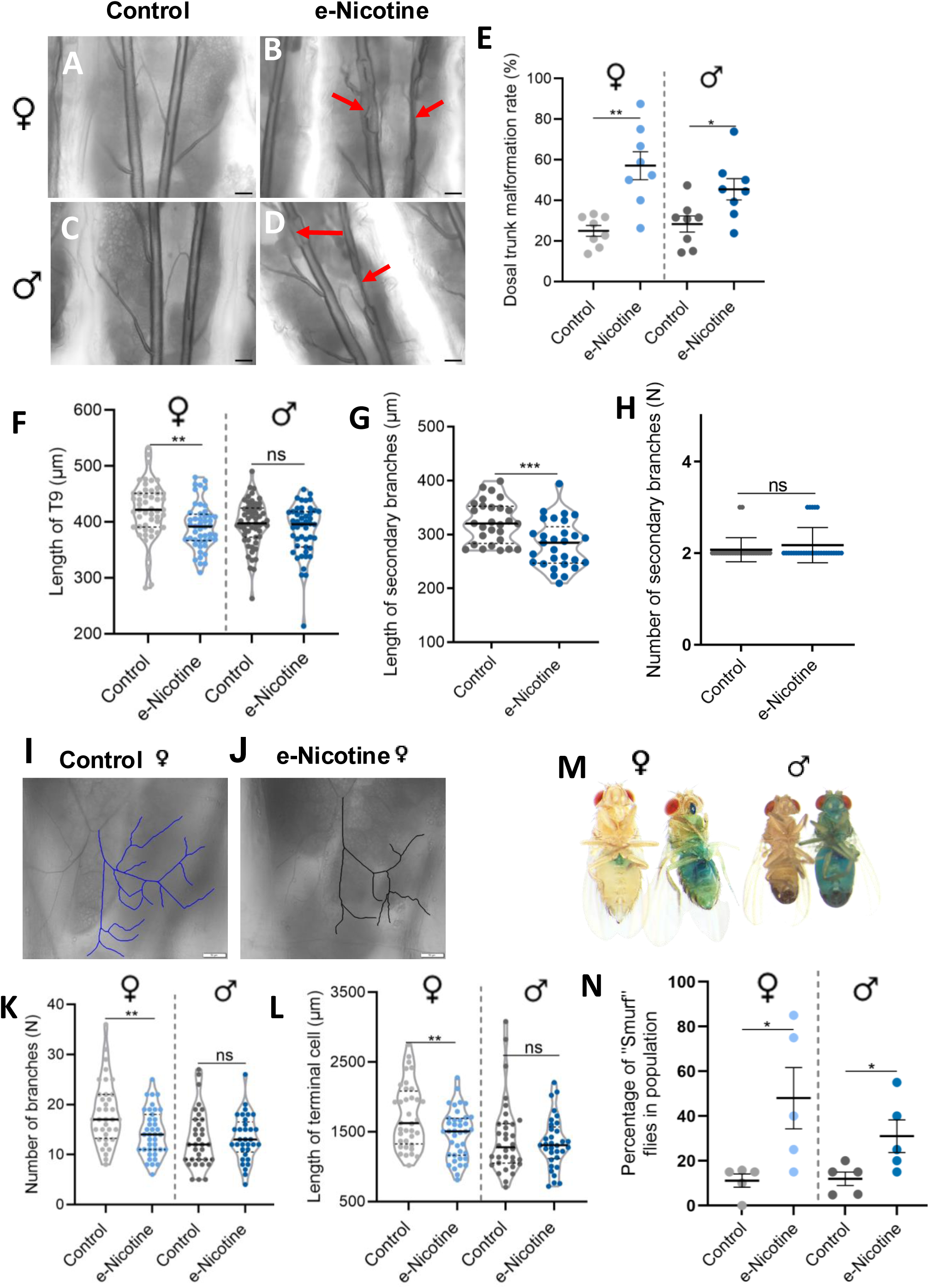
Preconception maternal e-nicotine exposure disrupts airway development and induces sex-specific structural defects in F_1_ offspring. **(A to D)** Representative bright field images of larval tracheal dorsal trunks in F_1_ offspring derived from control and e-nicotine exposed mothers. Female larvae are shown in A and B, and male larvae in C and D. Red arrows on the image mark tracheal narrowing in the offspring of e-nicotine exposed mothers. **(E)** Quantification of dorsal trunk deformation rate in F1 larvae from control and e-nicotine groups. Dorsal trunk deformation rate (%) in female offspring (24.95 ± 2.65% in control vs 57.09 ± 6.88% in larvae from e-nicotine exposed mothers; *P* = 0.0017). Dorsal trunk deformation rate in male offspring was 28.33 ± 3.92% in controls and 45.45 ± 5.26% in the e-nicotine group (*P* = 0.019). Each dot represents one independent biological replicate with 10-15 larvae per group. **(F)** Quantification of dorsal trunk length of the ninth tracheal segment (T9) in female (420.66 ± 7.20 µm in controls vs. 391.79 ± 5.89 µm in e-nicotine; n = 50 and 48, respectively; *P* = 0.0025), and male F1 larvae (396.28 ± 5.53 µm in controls vs. 385.64 ± 6.78 µm in e-nicotine; n = 56 and 47, respectively; *P* = 0.2268) from control and e-nicotine groups. (**G** and **H**) Quantification of the secondary tracheal branch length (G) and number (H) in the third tracheal segment (T3) in F1 larvae from control and e-nicotine groups with branch length (322.97 ± 7.36 µm in controls vs. 281.80 ± 8.46 µm in e-nicotine; n = 28 and 29, respectively; *P* = 0.0004) and branch number (2.07 ± 0.05 in controls vs. 2.17 ± 0.07 in e-nicotine; n = 28 and 29, respectively; *P* = 0.2508). (**I** and **J**) Representative bright field images of terminal tracheal cells in early third-instar F_1_ larvae from control and e-nicotine groups. Scale bars, 50 µm. (**K** and **L**) Quantification of terminal cell morphology in female and male F1 larvae from control and e-nicotine groups. Terminal branch number in females was 18.28 ± 1.06 in controls and 14.26 ± 0.74 in the e-nicotine group (n = 36 and 39, respectively; *P* = 0.0029), and in males was 13.12 ± 0.98 in controls and 13.30 ± 0.79 in the e-nicotine group (n = 34 and 33, respectively; *P* = 0.8836). Total terminal branch length in females was 1726.40 ± 77.19 µm in controls and 1460.46 ± 56.27 µm in the e-nicotine group (n = 36 and 39, respectively; *P* = 0.0070), and in males was 1412.41 ± 92.13 µm in controls and 1342.83 ± 63.77 µm in the e-nicotine group (n = 34 and 33, respectively; *P* = 0.5370). **(M)** Representative images of female and male “Smurf” flies displaying intestinal barrier dysfunction, indicated by systemic blue dye permeation. **(N)** Intestinal barrier integrity in adult F_1_ flies assessed using the Smurf assay. The proportion of flies exhibiting intestinal permeability was quantified in female and male offspring from control and e-nicotine groups. *P* values for comparisons between control and e-nicotine groups were 0.0307 in females and 0.0428 in males. Data in (E) were obtained from eight independent biological experiments, data in (F–H, K, L) from three independent biological experiments and five independent biological experiments for the Smurf assay (N). Data are presented as mean ± SEM. Statistical significance was determined using an unpaired two-tailed Student’s t test with Welch’s correction. Significance levels are indicated as follows: ns (*P* > 0.05), * (*P* < 0.05), **(*P* < 0.01), and *** (*P* < 0.001). Scale bars, 100 µm in (A–D) and 50 µm in (I, J).

Terminal cells in control offspring were extensively branched, whereas terminal cells in F_1_ female offspring from nicotine-exposed mothers appeared shorter and less branched (Fig. 1I, J). Quantitative analysis showed a significant reduction in total terminal branch length and branch number in female offspring, with no significant differences observed in males (Fig. 1K, L).

Following maternal preconception e-nicotine exposure, adult F_1_ flies of both sexes exhibited increased intestinal permeability, as indicated by the Smurf phenotype after blue dye feeding (Fig. 1M, N).

Notably, paternal e-nicotine exposure under the same conditions did not affect early developmental outcomes, including egg size, pupation, or hatching rates (Supplementary Fig. 2C–E), nor did it reproduce airway or terminal cell defects in offspring (Supplementary Fig. 2F–G), indicating that these phenotypes are specifically associated with maternal exposure.

These findings demonstrate that maternal preconception e-nicotine exposure selectively impairs terminal tracheal morphogenesis in female offspring, leading to shorter and less branched terminal cells, while male offspring remain largely unaffected. This sexual dimorphism suggests a female-specific vulnerability of distal airway development to maternal e-nicotine exposure.

### Maternal preconception e-nicotine induces epithelial thickening and airway remodelling in F_1_ larval tracheae

Given the structural abnormalities observed in the dorsal trunk and distal airways, we next examined whether maternal preconception e-nicotine exposure also affects the epithelial architecture of the tracheal system. We measured the epithelial thickness of the dorsal trunk (T9) and its main branches in offspring. Bright field micrographs showed a thin and evenly organized airway epithelium in control larvae (Fig. 2A), whereas offspring from e-nicotine-exposed mothers exhibited a visibly thicker epithelial layer (Fig. 2B). Quantitative analysis confirmed a consistent increase in epithelial thickness across the tracheal hierarchy (Fig. 2C–E). In the dorsal trunk (T9), epithelial thickness increased substantially after maternal e-nicotine exposure (Fig. 2C), which was also seen in the primary (Fig. 2D) and secondary branches (Fig. 2E). These results indicate that maternal preconception e-nicotine exposure induces a generalized epithelial thickening throughout the larval tracheal system, consistent with airway wall remodelling.

**Figure 2:**
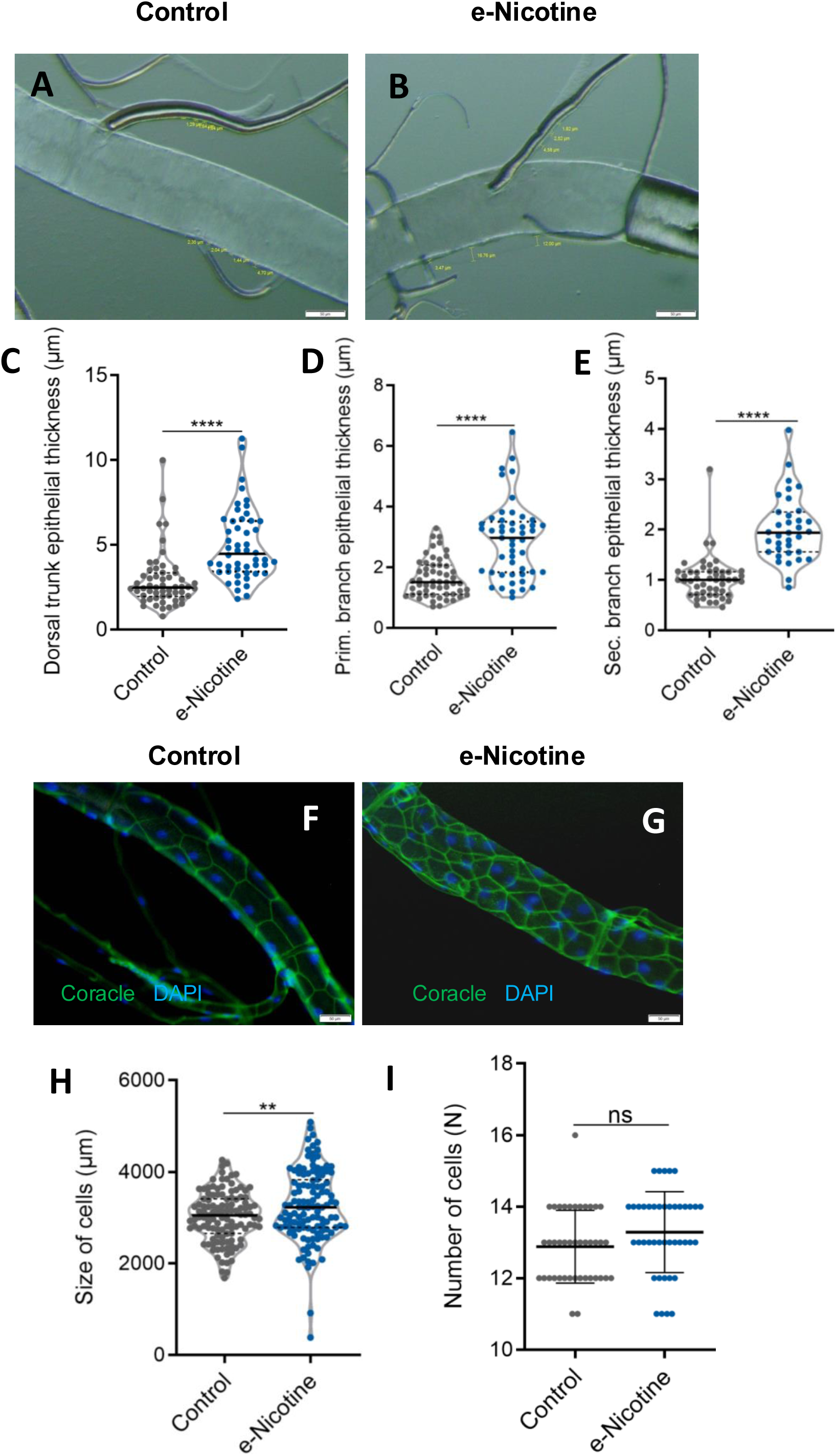
Maternal preconception e-nicotine exposure induces epithelial thickening and airway wall remodeling in larval tracheae. (**A** and **B**) Representative images of epithelial thickness in the tracheae of late third instar (L3) larvae from e-water (left) and e-nicotine (right) exposed mothers. (**C** to **E**) Quantification of epithelial thickness in the dorsal trunk (C) (2.88 ± 0.22 µm in controls vs. 4.97 ± 0.30 µm in e-nicotine, n = 55 and 50, respectively; *P* < 0.0001), primary branch (D) (1.64 ± 0.09 µm in controls vs. 2.89 ± 0.18 µm in e-nicotine, n = 55 and 49, respectively; *P* < 0.0001), and secondary branch (E) (1.02 ± 0.06 µm in controls vs. 2.01 ± 0.11 µm in e-nicotine, n = 49 and 35, respectively; *P* < 0.0001) of larval tracheae from control and maternal e-nicotine exposed offspring. (**F** and **G**) Representative immunofluorescence images of the tight junction protein Coracle in the ninth tracheal segment of the dorsal trunk, with nuclei counterstained with DAPI. (**H** and **I**) Quantification of epithelial cell area (3024 ± 50.80 µm² in controls vs. 3257 ± 71.01 µm² in e-nicotine, n = 115 and 120 cells, respectively; *P* = 0.0080) and cell number (12.88 ± 0.16 cells in controls vs. 13.29 ± 0.17 cells in e-nicotine, n = 42 and 42 cells, respectively; *P* = 0.0886) in the ninth segment of the dorsal trunk. Scale bars, 50 µm. Data were pooled from three independent biological experiments, with more than 10 larvae analyzed per experiment. Data are presented as mean ± SEM. Statistical analysis was performed using an unpaired two-tailed t test with Welch’s correction. ns, not significant. Significance levels are indicated as follows: ns (*P* > 0.05), **(*P* < 0.01), and **** (*P* < 0.0001).

In control offspring, Coracle, a tight junction protein lined the regular hexagonal pattern of airway epithelial cells, forming smooth and uniform epithelial walls (Fig. 2F). In contrast, offspring from mothers exposed to e-nicotine before conception exhibited irregularly shaped and enlarged epithelial cells, reflecting pronounced epithelial remodelling (Fig. 2G). The mean epithelial cell area was significantly increased in offspring from e-nicotine–exposed mothers (Fig. 2H), whereas the number of epithelial cells per T9 segment was not significantly different between groups (Fig. 2I), indicating that the observed airway wall thickening is attributable to epithelial cellular hypertrophy rather than hyperplasia.

### Developmental airway defects impair locomotor performance, hypoxia resilience, and survival in F_1_ offspring

We next assessed whether the structural airway and epithelial alterations observed in F_1_ offspring were associated with functional impairment by measuring locomotor performance and hypoxia resilience across larval and adult stages. In larvae, crawling distance was significantly reduced in both female and male F_1_ offspring from e-nicotine–exposed mothers (Fig. 3A). Consistent with this finding, tolerance to hypoxia (2–3% Ou) was significantly reduced in female larvae whereas no significant difference was observed in males (Fig. 3B).

**Figure 3:**
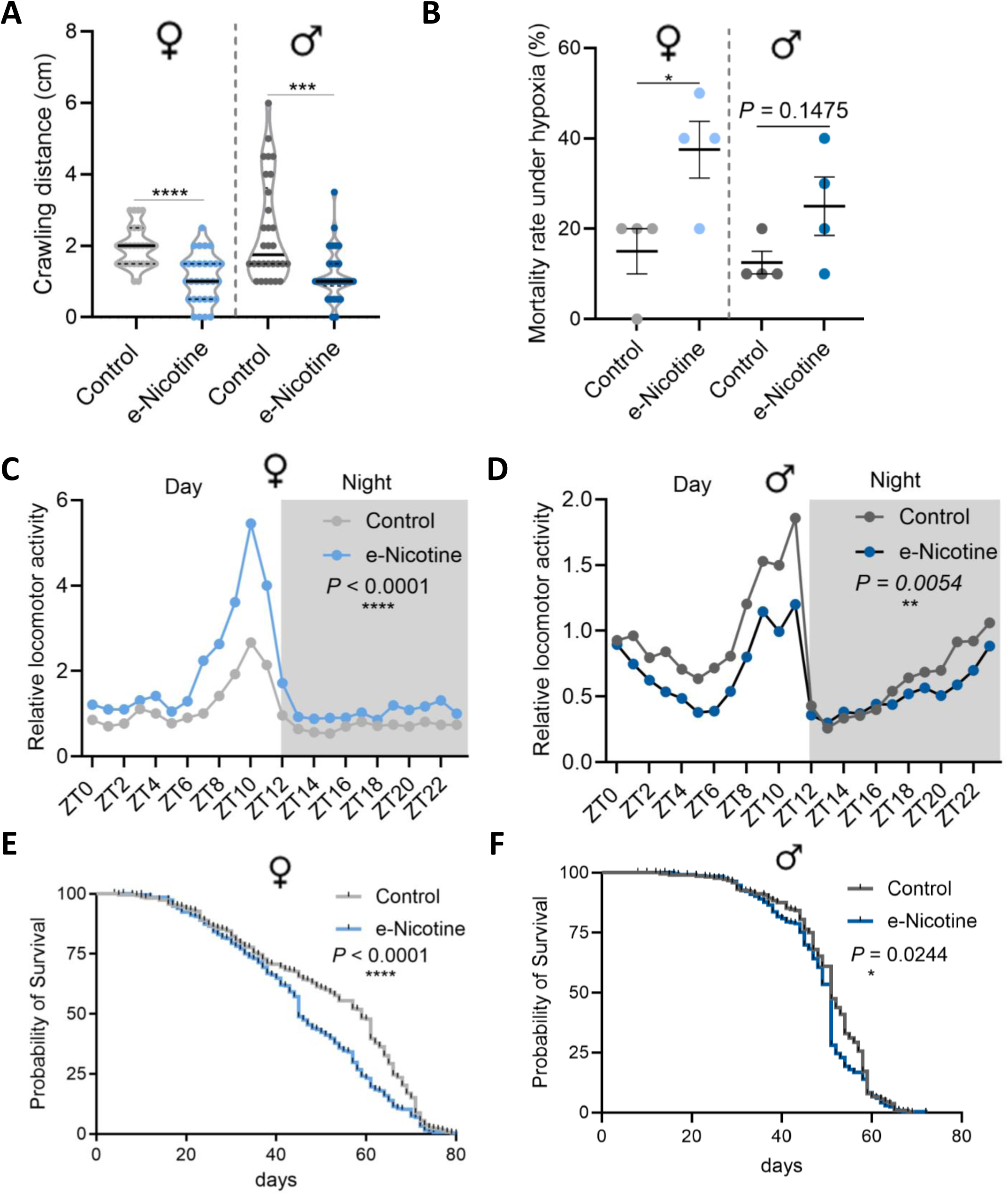
Airway developmental defects lead to functional decline and reduced lifespan in F_1_ offspring. **(A)** Crawling distance of female (1.97 ± 0.10 cm in controls vs. 1.08 ± 0.12 cm in e-nicotine; n = 30 per group; *P* < 0.0001) and male (2.40 ± 0.26 cm in controls vs. 1.18 ± 0.13 cm in e-nicotine; n = 10 per group; *P* = 0.0002) F_1_ larvae derived from control and e-nicotine exposure females. **(B)** Mortality of female (15.00 ± 5.00 % in controls vs. 37.50 ± 6.29 % in e-nicotine; *P* = 0.0329), and male (12.50 ± 2.50 % in controls vs. 25.00 ± 6.46 % in e-nicotine; n = 10 per group; *P* = 0.1475) F_1_ larvae following 3 h of severe hypoxia (2-3% OLJ). (**C** and **D**) Twenty-four-hour locomotor activity rhythms in adult female (C) and male (D) F_1_ flies derived from control and maternal e-nicotine exposed females. Activity was recorded across the 24-hour light dark cycle and plotted by Zeitgeber time (ZT). In females (C), time, treatment, and the time × treatment interaction significantly affected locomotor activity (n = 135 and 135, respectively; interaction, *P* < 0.0001; time, *P* < 0.0001; treatment, *P* < 0.0001). In males (D), time and treatment significantly affected locomotor activity (n = 135 and 135, respectively; time, *P* < 0.0001; treatment, *P* = 0.0054), whereas the time × treatment interaction was not significant (*P* = 0.0747). (**E** and **F**) Kaplan–Meier survival curves of adult female (E) and male (F) F_1_ flies derived from control and maternal e-nicotine exposed females. Median survival was 59 days in control females and 45 days in e-nicotine females (n = 284 and 286, respectively; *P* < 0.0001) and 51 days in both control and e-nicotine males (n = 265 and 268, respectively; *P* = 0.0244). Data were obtained from three independent biological experiments for crawling assays and locomotor activity analyses and from four independent biological experiments for hypoxia and lifespan analyses. Data are presented as mean ± SEM. Statistical significance was determined using an unpaired two-tailed Student’s t test with Welch’s correction for crawling distance, a two-sided Fisher’s exact test for mortality analysis, two-way repeated-measures ANOVA for locomotor activity, and the log-rank (Mantel–Cox) test for survival analysis. Significance levels are indicated as follows: ns (*P* > 0.05), * (*P* < 0.05), **(*P* < 0.01), and *** (*P* < 0.001), and **** (*P* < 0.0001).

Because locomotor impairment was observed in larvae from nicotine-exposed mothers, we next asked whether these functional alterations persist into adulthood. No significant differences in climbing ability were observed between offspring of e-nicotine exposed mothers and controls (Supplementary Fig. 3A). Monitoring locomotor activity patterns in adult flies over a 24-hour period revealed altered activity profiles in F_1_ offspring, characterized by increased activity in females and a mild reduction in males compared with controls (Fig. 3C, D). In contrast to the altered 24-hour activity patterns, baseline locomotor activity measured over a short 30-minute recording period was comparable between groups (Supplementary Fig. 3B). To further assess behavioral responsiveness, flies were exposed to a brief light stimulus. Following stimulation, locomotor response was significantly reduced in female F_1_ offspring, whereas no significant difference was observed in males (Supplementary Fig. 3C, D).

Additionally, lifespan was markedly reduced in female F_1_ offspring (Fig. 3E) and modestly reduced in males (Fig. 3F). Together, these findings indicate that maternal e-nicotine exposure is associated with altered behavioral performance and reduced physiological resilience in offspring.

### Airway transcriptomic profiling identifies Relish-centered immune repression with sex-specific network features

To determine whether maternal preconception e-nicotine exposure alters transcriptional programs in the next generation, we performed transcriptomic profiling of airways in female and male offspring and compared differentially expressed genes (DEGs) between sexes (Fig. 4A). Using an adjusted p-value threshold of padj < 0.05, females showed a relatively limited response, with 9 upregulated and 28 downregulated genes, whereas males displayed a broader response, with 48 upregulated and 81 downregulated genes. In females, the upregulated genes were dominated by members of the *Hsp70* family, including *Hsp70Aa, Hsp70Ab, Hsp70Ba, Hsp70Bb,* and *Hsp70Bc*, which encode canonical molecular chaperones involved in proteostasis, including protein folding and protection from proteotoxic stress (*35*) (Fig. 4A left). Notably, prior work in a *Drosophila* early-life cigarette-smoke exposure model had shown that smoke enters the larval airway system and induces *Hsp70* expression as part of an epithelial stress response (*33*). In contrast, the downregulated genes in females included several antimicrobial peptides, namely *CecA1, CecA2,* and *CecB*, together with *Relish* (*Rel/NF-*_κ_*B*), indicating suppression of a humoral immune program (*36*). Additional immune-associated or regulatory genes that were reduced included *PGRP-LF*, a negative regulator of Imd signaling that antagonizes *PGRP-LC* interactions (*37, 38*), and *Lk6*, a kinase linked to translational control through eIF4E phosphorylation and to growth and developmental regulation (*39*). Consistent with the volcano plot, heatmap visualization of the female DEG set showed clear separation of control and nicotine-treated samples, with both induced and repressed genes clustering coherently across biological replicates (Fig. 4B, left).

**Figure 4:**
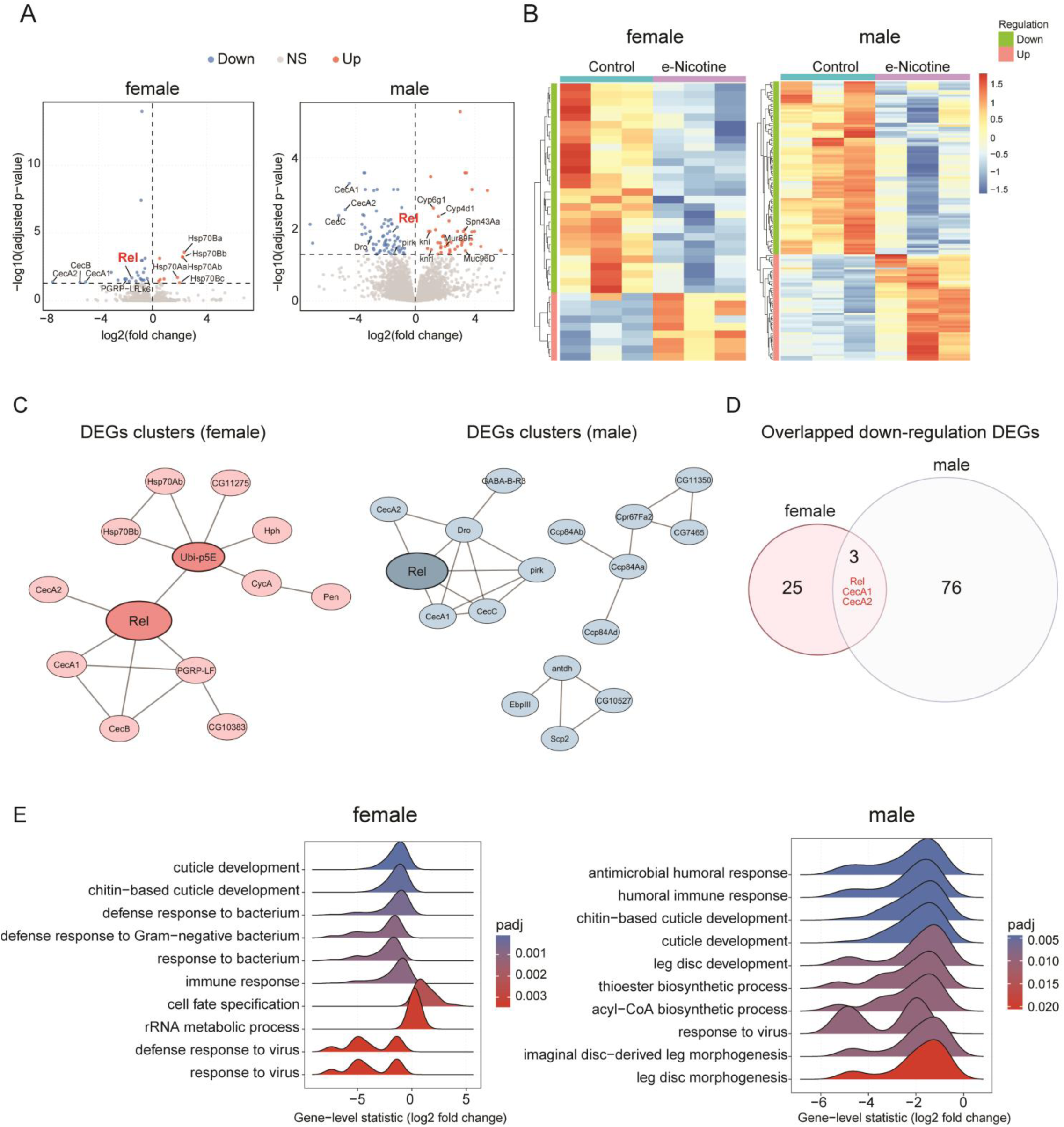
Airway transcriptomic profiling identifies Relish-centered immune repression with sex-specific network features. **(A)** Volcano plots showing differential gene expression in whole airways from female (left) and male (right) offspring comparing maternal preconception e-nicotine exposure versus control. Red dots indicate upregulated genes and blue dots indicate down regulated adjusted *P* value (padj value < 0.05). **(B)** Heatmaps of Differentially Expressed Genes (DEGs) in female (left) and male (right) airways from control and e-nicotine exposed groups. Expression values are shown as row-scaled normalized expression (z-scores). **(C)** Interaction-based clustering of female and male airway DEG sets. Networks were generated from STRING-derived interactions and visualized as gene interaction maps; nodes represent DEGs and edges represent functional associations. **(D)** Venn diagram showing overlap of downregulated DEGs between female and male airway datasets. **(E)** Gene set enrichment analysis (GSEA) of gene ontology (GO) Biological Process terms for female and male airway transcriptomes. Enrichment results are shown for the indicated gene sets and ranked by normalized enrichment score (NES) with multiple-comparisons–adjusted significance as displayed.

Males showed a stronger transcriptional response overall. Among the 81 downregulated genes, the most prominent included the antimicrobial peptides *CecC, CecA1,* and *CecA2* (Fig. 4A, right). Additional downregulated genes included *Dro* (*Drosomycin*), an antifungal antimicrobial peptide classically induced downstream of Toll signaling (*40*), and *pirk*, a well-established negative-feedback regulator of the Imd pathway that constrains pathway amplitude (*40, 41*). Like females, *Relish* (*Rel/NF-*_κ_*B*) was also downregulated in males (*42*), further supporting reduced Imd-associated transcriptional output in this condition. Heatmap analysis likewise showed clear segregation of control and nicotine-treated male samples, with coherent clustering of both upregulated and downregulated genes across replicates (Fig. 4B, right).

To further compare the transcriptional organization of female and male responses, we analyzed both DEG overlap and interaction-based clustering (Fig. 4C, D). Consistent with the limited overlap between sexes, no shared genes were detected among upregulated DEGs, whereas only three genes overlapped among downregulated DEGs, namely *Relish* (*Rel/NF-*_κ_*B*), *CecA1,* and *CecA2*. This indicates that female and male offspring largely engaged distinct transcriptional programs, while the small, shared component converged on a common immune core centered on *Relish*. Consistent with this, interaction-based clustering showed that the female DEG network was relatively compact and centered on *Relish*, consistent with a small and coherent immune-stress module. By contrast, the male DEG network was more distributed and resolved into several subnetworks, including a prominent *Relish*-centered immune cluster together with additional modules linked to cuticle-associated and metabolic functions. Thus, while the female response appeared more focused, the male response extended across multiple functional modules.

Finally, we used gene set enrichment analysis (GSEA) with Gene Ontology Biological Process terms to identify broader biological programs associated with these transcriptional changes (Fig. 4E). In both females and males, the top 10 negatively enriched GO terms were predominantly related to immune processes and cuticle development, consistent with an overall reduction in these programs following maternal preconception e-nicotine exposure.

### NF-_κ_B/Relish suppression phenocopies e-nicotine-induced airway and behavioral defects

Since Relish (Rel/NF-κB), a core transcription factor in the Imd pathway controlling AMP programs (*43, 44*), was downregulated in both males and females, we next investigated whether suppression of Rel/NF-κB signalling contributes to maternally e-nicotine induced phenotypes. Tracheal specific knockdown of Relish using *Btl-Gal4>UAS-Relish-RNAi* resulted in reduced body size at both larval and adult stages (Fig. 5A–B), consistent with previously reported growth defects in offspring following maternal e-nicotine exposure (*8*). In line with the phenotypes observed after maternal e-nicotine exposure (Fig. 1A-E), Relish knockdown led to increased airway abnormalities (Fig. 5C).

**Figure 5:**
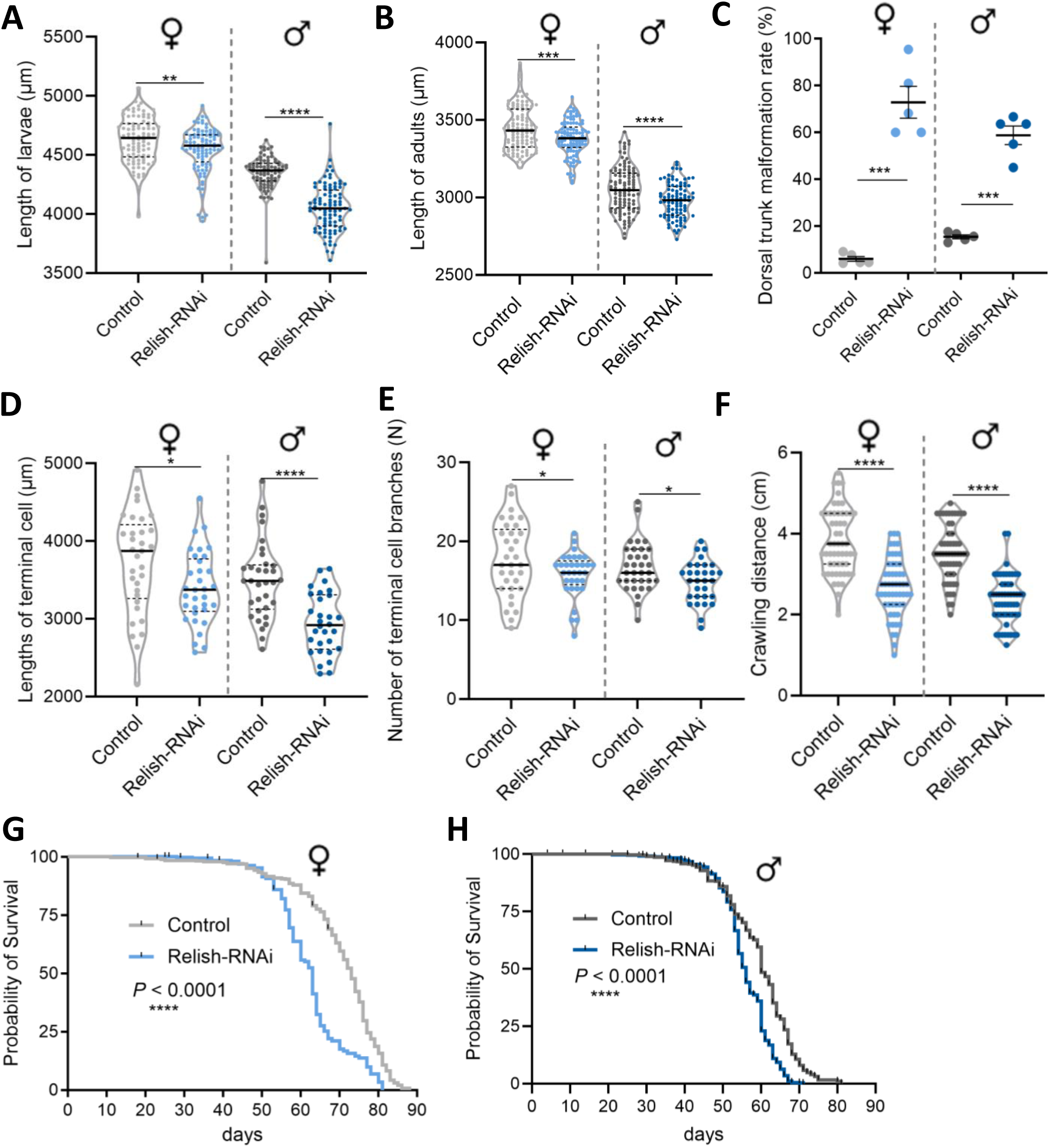
Trachea-specific Relish knockdown phenocopies maternal e-nicotine-induced airway and behavioral phenotypes in F_1_ offspring. **(A)** Body length of larvae in female (4626.91 ± 19.87 µm in controls vs. 4543.47 ± 19.41 µm in Relish-RNAi; *P* = 0.003) and male (4351.79 ± 14.73 µm in controls vs. 4052.40 ± 19.91 µm in Relish-RNAi; *P* < 0.0001) flies with trachea-specific Relish knockdown (Relish-RNAi) compared with control flies. **(B)** Body length of adults, in female flies (3444 ± 17 µm in controls vs. 3381 ± 17 µm in Relish-RNAi; n = 107 and 110, respectively; *P* = 0.0003) and in male flies (3053 ± 18 µm in controls vs. 2979 ± 18 µm in Relish-RNAi; n = 108 and 96, respectively; *P* < 0.0001). **(C)** Percentage of larvae exhibiting dorsal trunk malformations in female and male flies following trachea-specific Relish knockdown. Values in females were 6.01 ± 6.89% in controls vs. 72.92 ± 6.89% in Relish-RNAi (*P* = 0.0005) and in males were 15.46 ± 4.05% in controls vs. 58.79 ± 4.05% in Relish-RNAi (*P* = 0.0003). (**D** and **E**) Terminal cell morphology in late third-instar larvae following terminal cell–specific Relish knockdown. Terminal cell length (D) in females was 3727.51 ± 113.58 µm in controls and 3395.79 ± 81.31 µm in Relish-RNAi (*P* = 0.0209; n = 33 and 33, respectively), and in males was 3494.42 ± 86.03 µm in controls and 2937.74 ± 70.55 µm in Relish-RNAi (*P* < 0.0001; n = 33 and 30, respectively). The number of terminal cell branches (E) in females was 17.73 ± 0.82 in controls and 15.79 ± 0.49 in Relish-RNAi (*P* = 0.0475; n = 33 and 33, respectively), and in males was 16.73 ± 0.54 in controls and 14.90 ± 0.48 in Relish-RNAi (*P* = 0.0145; n = 33 and 30, respectively). **(F)** Crawling distance of female (3.80 ± 0.11 cm in controls vs. 2.67 ± 0.10 cm in Relish-RNAi; n = 55 per group; *P* < 0.0001) and male larvae (3.53 ± 0.09 cm in controls vs. 2.37 ± 0.08 cm in Relish-RNAi; n = 55 per group; *P* < 0.0001) with trachea-specific Relish knockdown compared with control larvae. (**G** and **H**) Kaplan–Meier survival analysis of female (G) and male (H) flies following trachea-specific Relish knockdown. Median lifespan was reduced from 73 days to 63 days in females and from 60 days to 56 days in males following Relish knockdown (*P* < 0.0001 for both comparisons; n = 385, 310, 380, and 348, respectively). Flies in panels A–C and G–H were generated using the trachea-specific driver *btl-Gal4, UAS-GFP* crossed to either *UAS-Rel-RNAi* or the corresponding RNAi control line, whereas flies in panels D–F were generated using the terminal cell–specific driver *DSRF-Gal4, UAS-GFP* crossed to the same lines. Data were obtained from four independent biological experiments for body length, locomotor, and lifespan assays (panels A, B, F, G, and H), five independent biological experiments for dorsal trunk malformation analysis (panel C), and three independent biological experiments for terminal cell measurements (panels D and E). Data are presented as mean ± SEM. Statistical significance was determined using an unpaired two-tailed Student’s t test with Welch’s correction. Significance levels are indicated as follows: * (*P* < 0.05), **(*P* < 0.01), *** (*P* < 0.001), and **** (*P* < 0.0001).

To further define the cellular basis of these developmental defects, we performed terminal cell–specific knockdown of Relish using *DSRF-Gal4 > UAS-Relish-RNAi*. Loss of Relish function significantly reduced terminal cell length and branching complexity in late third-instar larvae (Fig. 5D–E), phenocopying the terminal cell defects observed following maternal e-nicotine exposure (Fig. 1I–L).

Consistent with these structural abnormalities, larvae with Relish knockdown displayed significantly reduced locomotor performance (Fig. 5F), similar to the behavioral deficits observed in nicotine-exposed offspring (Fig. 3A). Moreover, trachea-specific suppression of Relish resulted in a pronounced reduction in lifespan in both female and male flies (Fig. 5G–H), paralleling the shortened lifespan observed in offspring following maternal e-nicotine exposure (Fig. 3E-F).

Together, these findings demonstrate that suppression of NF-κB/Relish signalling is sufficient to phenocopy the developmental, airway, behavioral, and survival defects induced by maternal e-nicotine exposure.

### Preconception e-nicotine exposure reduces proliferation of F_1_ airway epithelial progenitor cells

Given the observed impairments in locomotor performance, behavioral responsiveness, and lifespan in adult F_1_ offspring, we next examined whether these functional deficits originate from altered airway development by assessing the progenitor compartment of developing tracheae. These tracheoblasts reside in larval tracheal segments Tr4 and Tr5, where they form specialized niches that regulate progenitor maintenance and activation (*32, 45, 46*).

Fluorescence micrographs of the Tr5 tracheal progenitor niche stained with DAPI and shown in grayscale to reflect raw signal intensity revealed large, well-defined progenitor niche domains in control female offspring (Fig. 6A), whereas female offspring from nicotine-exposed mothers displayed visibly smaller niches (Fig. 6B). A similar reduction was observed in male offspring, as illustrated in the Tr5 niche of control and e-nicotine groups (Fig. 6C, D). Quantitative analysis confirmed a significant reduction in Tr5 niche area in both female and male offspring following maternal preconception nicotine exposure (Fig. 6E). A similar reduction was also observed in the Tr4 progenitor niche (Supplementary Fig. 4).

**Figure 6:**
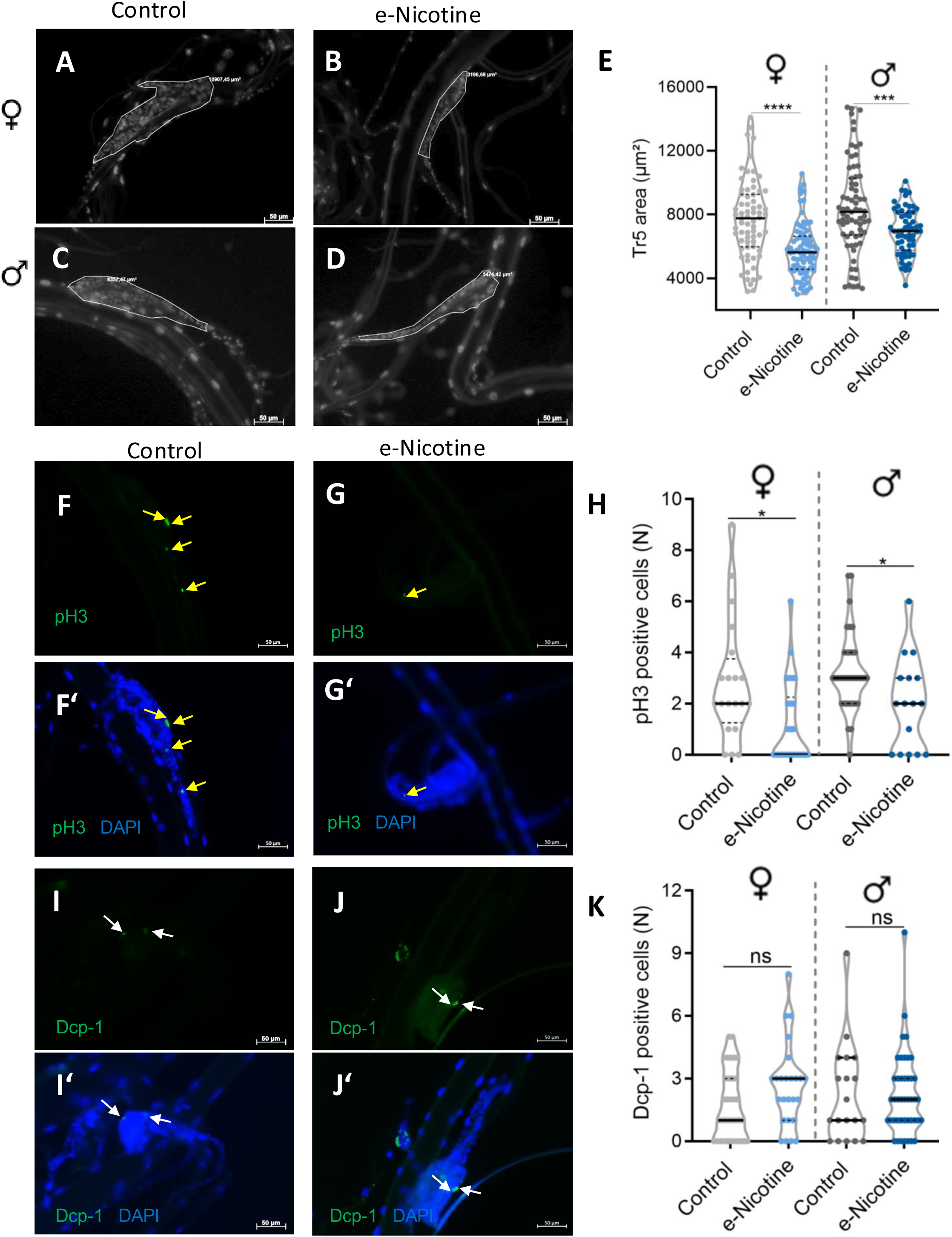
Preconception e-nicotine exposure impairs tracheal progenitor proliferation without inducing apoptosis in F_1_ larvae. (**A** to **D**) Representative fluorescence images of the Tr5 tracheal stem cell niche in female (A, B) and male (C, D) F1 larvae derived from control and maternal e-nicotine exposed groups. Stem cell nuclei were visualized using DAPI staining to define the boundaries of the Tr5 stem cell niche. (**E**) Quantification of Tr5 stem cell niche area in female (7686.56 ± 295.59 µm² in controls vs. 5780.52 ± 196.72 µm² in e-nicotine, n = 69 and 75, respectively; *P* < 0.0001) and male F_1_ larvae (8425.45 ± 342.69 µm² in controls vs. 6940.24 ± 188.04 µm² in e-nicotine, n = 71 and 60, respectively; *P* = 0.0002). (**F** to **H**) Representative immunofluorescence images of proliferating cells in the Tr5 stem cell niche detected by phospho-histone H3 (pH3) staining. Yellow arrows indicate pH3-positive cells. Representative images are from females. Quantification of pH3-positive cells in the Tr5 stem cell niche is shown in (H), with *P* values of 0.0115 in females (n = 20 and 26) and 0.0253 in males (n = 35 and 16). (**I** to **K**) Representative immunofluorescence images of apoptotic cells in the Tr5 stem cell niche detected by cleaved Dcp-1 staining. White arrows indicate Dcp-1 positive apoptotic cells. Representative images are from female larvae. Quantification of Dcp-1-positive cells in the Tr5 stem cell niche is shown in (K), with *P* values of 0.0650 in females (n = 35 and 25) and 0.9715 in males (n = 19 and 43). Data were pooled from three independent biological experiments, with more than 15 larvae dissected per group in each experiment. Data are presented as mean ± SEM. Statistical analysis was performed using an unpaired two-tailed t test with Welch’s correction. Scale bars, 50 µm. Significance levels are indicated as follows: ns (*P* > 0.05), * (*P* < 0.05), *** (*P* < 0.001), and **** (*P* < 0.0001).

As it was unclear what caused the reduction in tracheoblast niche area, we assessed mitotic and apoptotic activity in the Tr5 tracheoblast niche by immunofluorescence staining for phosphorylated histone H3 (pH3) (*47*) and Death caspase-1 (Dcp-1) (*48*), respectively.

Fluorescence images showed abundant pH3-positive nuclei in control offspring (Fig. 6F, F′), whereas the number of dividing cells was lower in offspring from e-nicotine mothers (Fig. 6G, G′). Quantitative analysis confirmed the significant reduction in mitotic activity (Fig. 6H). By contrast, staining for the apoptotic marker Dcp-1 revealed no significant difference between groups (Fig. 6I–K).

Together, these findings demonstrate that maternal preconception exposure to e-nicotine compromises the proliferative capacity of tracheal progenitor niches, leading to a reduction in the overall size of tracheoblast niches.

### Tracheal stem-cell niche transcriptomics reveals sex-specific immune repression and airway remodeling programs

The reduction in airway stem-cell niche size and progenitor proliferation prompted transcriptomic profiling of Tr4/Tr5 niches in late third-instar larvae following maternal preconception e-nicotine exposure (Fig. 7A). Differential expression analysis revealed distinct transcriptional responses between female and male niches (Fig. 7B). Volcano plot analysis revealed a striking sex-specific bias in transcriptional responsiveness. Females exhibited a limited signature with only one significantly downregulated transcript, mt:ND6, and a modest set of upregulated genes (Fig. 7B left). By contrast, males showed a substantially broader response with extensive gene repression and induction (Fig. 7B right).

**Figure 7:**
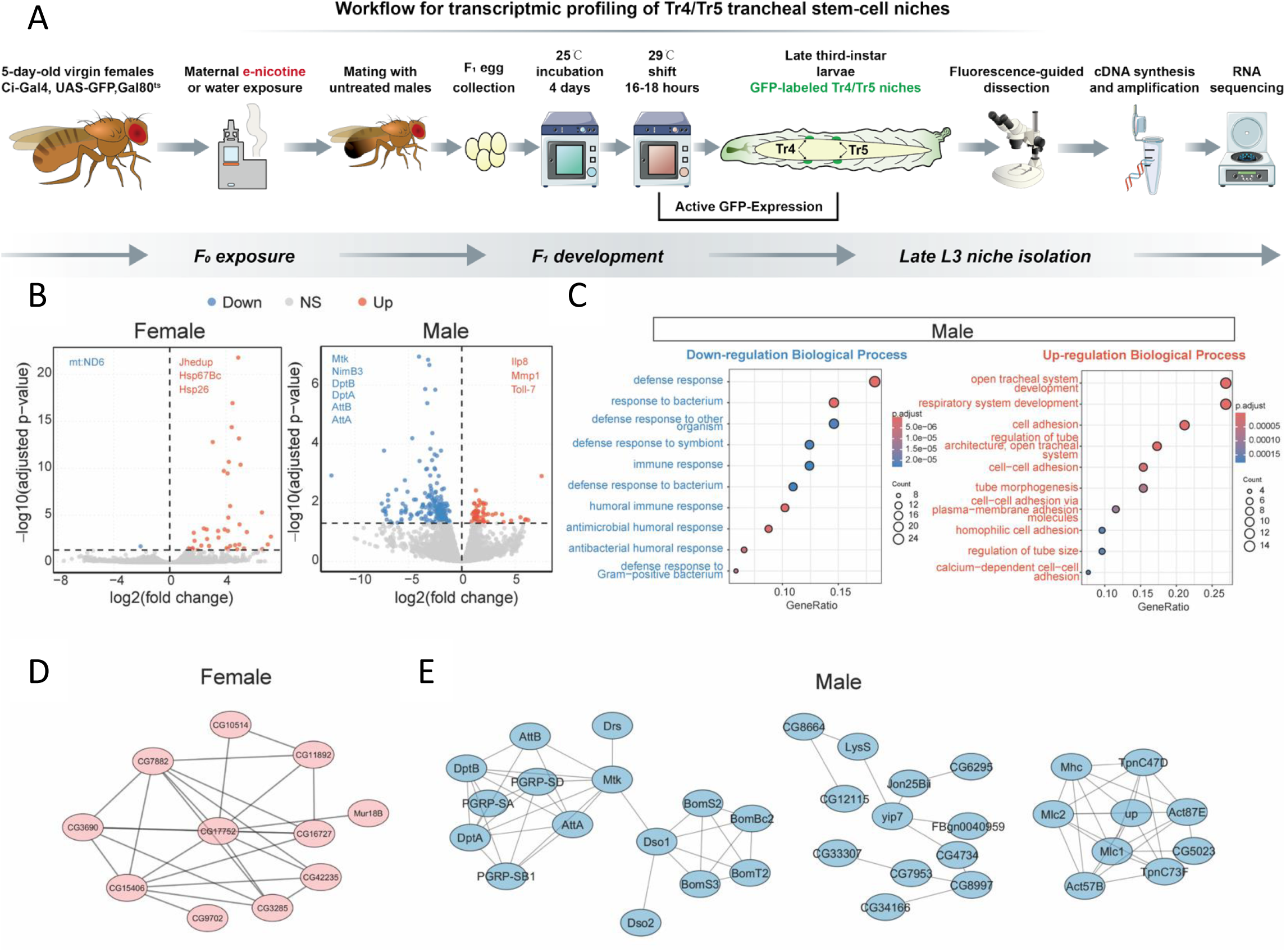
Sex-specific transcriptional and network features in the larval Tr4/Tr5 tracheal stem-cell niche following maternal preconception e-nicotine exposure. (**A**) Experimental workflow for transcriptomic profiling of Tr4/Tr5 tracheal stem-cell niches. Five-day-old virgin *Ci-Gal4, UAS-GFP; Gal80ts* females were exposed to e-nicotine or water (vehicle control) prior to mating with untreated males. F_1_ eggs were collected and larvae were reared at 25°C for 4 days, followed by transfer to 29°C for 16–18 h to induce GFP labelling of Tr4/Tr5 stem-cell niches through Gal80ts inactivation. GFP-positive niches were isolated under a fluorescence stereomicroscope and processed for RNA extraction, cDNA synthesis, amplification, and sequencing. (**B**) Volcano plots showing differential gene expression in Tr4/Tr5 tracheal stem-cell niche samples from female and male offspring comparing maternal e-nicotine versus control conditions. Each point represents one gene; dashed lines indicate the thresholds used to define DEGs. Red dots indicate upregulated genes, and blue dots indicate down regulated (adjusted *P* value, padj < 0.05; absolute log2 fold change, |log2FC| > 1). (**C**) GO Biological Process enrichment analysis of male DEGs, performed separately for upregulated and downregulated gene sets. Bubble size indicates the number of genes annotated to each term, while colour represents the FDR for enrichment significance. (**D** and **E**) STRING protein-protein interaction networks of female (D) and male (E) DEGs using high-confidence interactions (STRING score ≥ 0.7).

In females, the induced gene set was dominated by stress- and metabolism-associated factors. Notably, small heat-shock proteins (Hsp26 and Hsp67Bc) were strongly upregulated (Fig. 7B), consistent with enhanced proteostasis and cellular stress handling in the niche. In addition, several enzymes and transporters linked to metabolic regulation and xenobiotic handling were increased, including the esterase-related genes Jhe and Jhedup, the detoxification enzyme GstT3, and the transporter Oatp58Da. Together, these changes point to a focused stress–metabolic adaptation program in the female niche rather than broad pathway repression.

Male niche transcriptional changes were organized into two majors, functionally opposing trends. First, the downregulated genes were strongly enriched for innate immune effectors, including multiple antimicrobial peptides (e.g., *DptA/DptB*, *AttA/AttB*, *Mtk*, and *Drs*) and upstream immune recognition components (*PGRP-SA*, *PGRP-SD*, and *PGRP-SB1*). Gene Ontology enrichment of male downregulated DEGs confirmed that immune-related biological processes were the dominant suppressed program (Fig. 7C). Second, the male upregulated DEGs preferentially mapped to airway structural and developmental programs. GO enrichment highlighted terms such as open tracheal system development, respiratory system development, and cell adhesion, consistent with engagement of airway remodeling pathways in response to maternal exposure (Fig. 7C). Representative induced genes included the damage-associated signal Ilp8 and extracellular matrix remodeling factors (Mmp1 and AdamTS-A), together with multiple genes linked to epithelial architecture and adhesion.

Network-level analysis further supported distinct, sex-specific organization of niche DEGs. In females, STRING interaction mapping resolved into a single compact cluster composed largely of poorly annotated CG genes, with CG17752 emerging as a central node connecting multiple induced partners, consistent with a focused and highly interconnected female niche response (Fig. 7D). By contrast, males revealed a modular architecture with three major interaction clusters: a densely connected humoral immune effector module linking antimicrobial peptides and PGRP components (e.g., *DptA/DptB*, *AttA/AttB*, *Mtk, Drs*, and *PGRP-SA/SD/SB1*), a Bom/Dso-associated subnetwork, and a structural/cytoskeletal module containing actin–myosin and troponin-associated genes (e.g., *Act57B*, *Act87E*, *Mhc*, *Mlc1/2*, and *TpnC47D/TpnC73F*) (Fig. 7E). Collectively, these interaction patterns indicate that maternal preconception e-nicotine exposure imprints the tracheal stem-cell niche in a sex-dependent manner, with females showing a more limited DEG-associated interaction module and males exhibiting broader, multi-module reprogramming that spans immune repression and structural remodeling-associated networks.

### Maternal preconception e-nicotine exposure propagates developmental and airway phenotypes across generations with progressive attenuation

Finally, we assessed whether developmental alterations in F_1_ induced by maternal e-nicotine persist into subsequent generations. Successive generations were generated using a controlled breeding strategy in which exposed F_׈_ females were crossed with untreated males, and subsequent generations were propagated using non-exposed females from each lineage (Fig. 8A). In the F_2_ generation, offspring derived from F_1_ parents born to nicotine-exposed mothers exhibited a significant reduction in egg length compared with F_2_ offspring from control lineages (Fig. 8B), demonstrating that developmental effects persist into the next generation despite the absence of direct exposure. This phenotype persisted through larval development, as F_2_ larvae from exposed lineages showed a significant reduction in body length compared with controls (Fig. 8H). Furthermore, the reduction in body size persisted into adulthood, as F_2_ generation flies displayed significantly shorter adult body length compared with controls (Fig. 8E), suggesting that the growth impairment observed during early development persisted later in life. As seen for the parental F_1_ generation from e-nicotine mothers, F_2_ larvae had a significant decrease in crawling distance (Fig. 8I), indicating impaired functional consequence during larval stages. Of note, these developmental deficits also extended to the next generation, as F_3_ embryos also exhibited reduced egg length compared with controls (Fig. 8C), demonstrating that the phenotype is maintained across generations. However, this phenotype was no longer detectable at later stages, as adult body length in F_3_ flies did not differ from controls in either sex (Fig. 8F), and no difference was observed in F_4_ adults (Fig. 8D, G).

**Figure 8:**
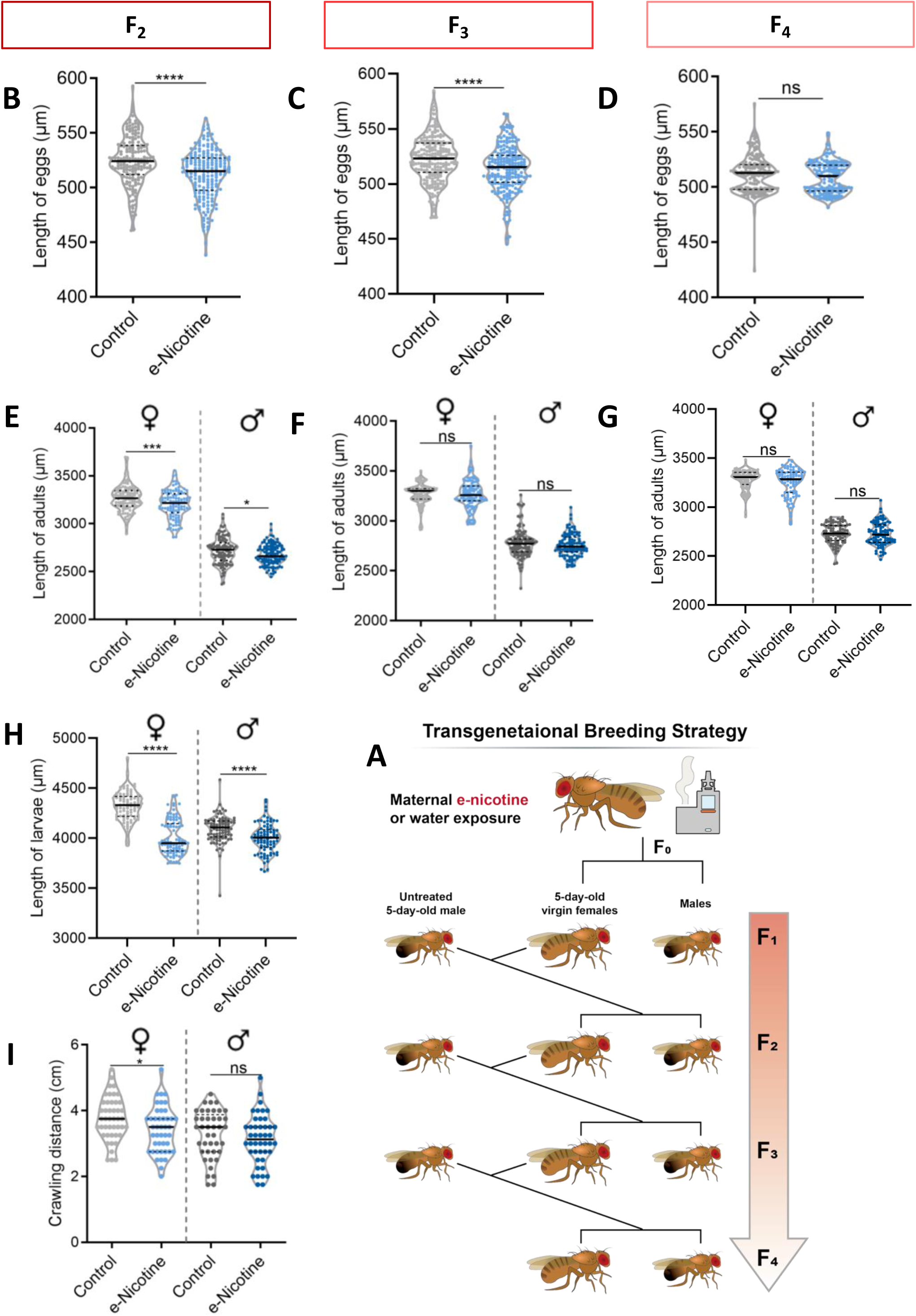
Maternal preconception e-nicotine exposure propagates developmental and airway phenotypes across generations with progressive attenuation. (**A**) Transgenerational breeding strategy used to generate F_2_–F_4_ progeny from control and maternal e-nicotine lineages. (**B**, **C**, **D**) Egg length in F_2_ (B), F_3_ (C), and F_4_ (D) progeny from control and maternal e-nicotine exposed lineages. Values were 524.60 ± 1.71 µm and 512.16 ± 1.60 µm in F_2_ (n = 169 and 193; *P* < 0.0001), 522.79 ± 1.42 µm and 513.92 ± 1.43 µm in F_3_ (n = 209 and 207; *P* < 0.0001), and 511.21 ± 1.26 µm and 508.75 ± 1.08 µm in F_4_ (n = 182 and 163; *P* = 0.1384). (**E**, **F**, **G**, **H**) Body length of F_2_ larvae (H), F_2_ adults (E), F_3_ adults (F), and F_4_ adults (G) in female and male progeny from control and maternal e-nicotine–exposed lineages. In F_2_ larvae, body length in females was 4321.11 ± 15.10 µm in controls and 4007.21 ± 18.14 µm in the e-nicotine lineage (n = 92 and 90; *P* < 0.0001), and in males was 4092.19 ± 14.12 µm and 3999.89 ± 15.62 µm (n = 91 and 88; *P* < 0.0001). In F_2_ adults, body length in females was 3267.00 ± 12.65 µm and 3202.05 ± 14.12 µm (n = 113 and 105; *P* = 0.0007), and in males was 2716.51 ± 13.27 µm and 2677.05 ± 9.48 µm (n = 109 and 122; *P* = 0.0164). In F_3_ adults, body length in females was 3267.68 ± 10.08 µm and 3263.25 ± 13.58 µm (n = 122 and 111; *P* = 0.7936), and in males was 2771.41 ± 12.52 µm and 2755.94 ± 10.75 µm (n = 112 and 108; *P* = 0.3495). In F_4_ adults, body length in females was 3279.89 ± 9.80 µm and 3251.66 ± 14.85 µm (n = 105 and 92; *P* = 0.1146), and in males was 2730.38 ± 10.19 µm and 2724.36 ± 11.49 µm (n = 99 and 101; *P* = 0.6952). (**I**) Crawling distance in F_2_ female and male larvae from control and maternal e-nicotine exposed lineages. Crawling distance in females was 3.76 ± 0.10 cm in controls and 3.41 ± 0.10 cm in the e-nicotine lineage (n = 45 and 44; *P* = 0.0194), and in males was 3.28 ± 0.11 cm and 3.13 ± 0.12 cm (n = 41 and 44; *P* = 0.3349). Data were obtained from three independent biological experiments. Data are presented as mean ± SEM. Statistical significance was determined using an unpaired two-tailed Student’s t test with Welch’s correction. Significance levels are indicated as follows: ns (*P* > 0.05), * (*P* < 0.05), *** (*P* < 0.001), and **** (*P* < 0.0001).

**Figure 9:**
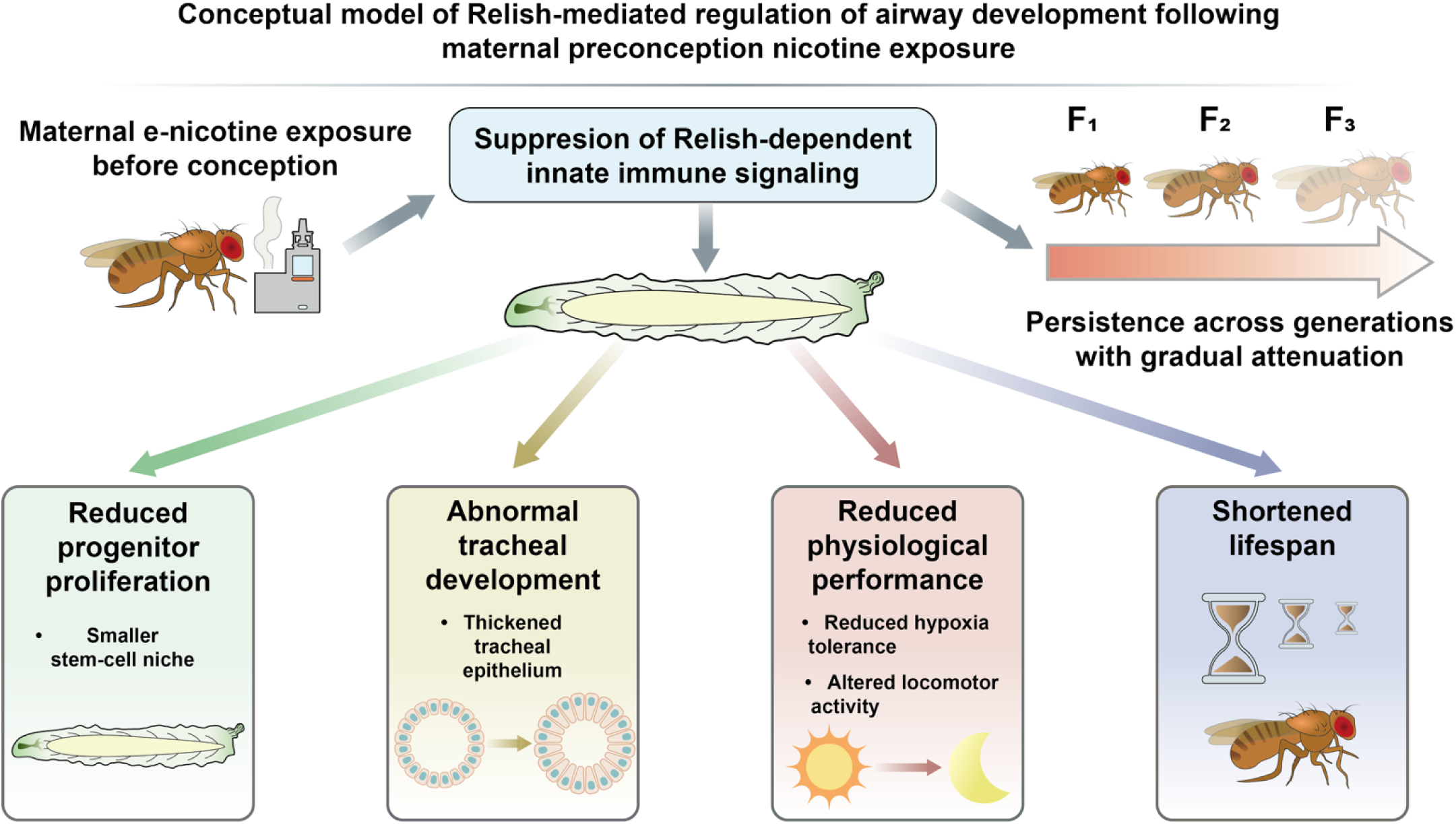
Conceptual model of maternal preconception e-nicotine–induced airway niche reprogramming via NF-κB/Relish signaling. Preconception nicotine exposure suppresses Relish-dependent innate immune signaling in the airway epithelium, leading to reduced progenitor proliferation, abnormal airway development, impaired physiological performance, and shortened lifespan. These developmental and functional defects persist across generations with gradual attenuation, highlighting Relish signaling as a central regulator of airway developmental programming.

## Discussion

The present study demonstrates that maternal exposure to vaporized nicotine prior to conception is sufficient to disrupt normal airway morphogenesis and compromise physiological function in the next generation. We show for the first time that this is linked to altered airway and progenitor regulatory programs with reduced progenitor cell expansion. Mechanistically, our data implicate Relish/NF-κB signalling as a key pathway contributing to the observed developmental abnormalities. While previous work has predominantly focused on gestational exposure, our results substantiate the preconception period as an important parental window during which e-nicotine can influence developmental trajectories and physiological resilience in offspring.

Previous human and experimental studies have shown that maternal smoking or prenatal nicotine exposure compromises fetal lung development and reduces pulmonary function in offspring, supporting the idea that early airway defects can have lasting physiological consequences (*16, 18, 20*). Maternal e-nicotine exposure also led to increased epithelial thickness throughout the airway tree in F_1_. This increase was primarily linked to enlargement of epithelial cells, showing cellular hypertrophy rather than hyperplasia. Epithelial thickening and altered cellular organization are well-established features of airway remodelling following injury or environmental exposure (*49, 50*). Consistent with this pattern, studies in mammalian systems have shown that prenatal nicotine or cigarette smoke exposure leads to airway wall thickening and impaired epithelial differentiation in the developing lung (*16, 51*). Together, these findings suggest that disturbances in airway development can be accompanied by structural remodelling of the airway epithelium, which may contribute to long-term impairment of airway function (*52*).

Developmental airway abnormalities in offspring from e-nicotine mothers were associated with reduced larval physiological performance, as evidenced by decreased locomotor activity. Although tissue oxygenation was not directly measured in the present study, the reduced tolerance to hypoxic stress observed in exposed offspring is consistent with the functional consequences of impaired airway structure. and lower tolerance to hypoxic stress. These findings indicate that structural airway defects translate into measurable functional limitations rather than being purely anatomical changes. These observations suggest that disturbances in airway development can affect the ability of the organism to cope with environmental stress. Similar functional impairments have been reported in mammalian studies of prenatal nicotine or maternal smoking exposure, where offspring show reduced respiratory performance and increased vulnerability to hypoxia-related stress (*51, 53, 54*). Taken together, these findings support the view that early disruption of airway development can lead to lasting functional consequences in nicotine-exposed offspring.

Functional abnormalities in offspring of e-nicotine-exposed mothers persisted into adulthood. In addition to reduced lifespan, adult flies displayed disrupted locomotor activity patterns and diminished responsiveness to external stimuli, indicating compromised physiological resilience. Altered daily activity rhythms suggest dysregulation of systemic homeostasis, while reduced responses to light stimulation point to impaired stress adaptability rather than a primary deficit in baseline locomotion (*55*). Together, these findings demonstrate that maternal e-nicotine induces sustained physiological consequences that persist beyond early developmental stages (*56, 57*). The combination of altered circadian activity, reduced stress responsiveness, and shortened lifespan highlights impaired physiological robustness in adulthood, and underlines the long-term impact of maternal nicotine exposure on organismal health.

Transcriptomic profiling of larval tracheae revealed suppression of innate immune signalling pathways following maternal preconception e-nicotine exposure, with Relish (NF-κB) identified as a shared downregulated transcription factor across sexes. NF-κB signalling regulates epithelial defence and stress responses in *Drosophila* (*58*) and is also activated in airway epithelial cells of smokers and in experimental nicotine exposure models (*59, 60*), indicating its responsiveness to tobacco-related stimuli. Reduced *Relish* expression therefore suggests disruption of processes required for normal airway development. Functional experiments support this interpretation: trachea-specific *Relish* knockdown reproduced key phenotypes observed after maternal e-nicotine exposure, including reduced body size, airway deformation, impaired terminal branching, and decreased locomotor performance. Together, these findings identify NF-κB/Relish signalling as a candidate pathway contributing to e-nicotine-induced defects in airway development and physiological function in offspring.

Previous studies of prenatal nicotine or cigarette smoke exposure consistently reported persistent abnormalities in lung structure and function in offspring, but the underlying developmental processes remain poorly defined (*19, 32*). Here, we provide evidence that impaired expansion of airway progenitor cells may link maternal e-nicotine exposure to long-term airway dysfunction. Rather than reflecting irreversible tissue damage, the reduced progenitor cell niche size suggests a failure of normal developmental renewal during airway growth. This extends our previous work showing lasting growth defects in offspring (*8*), and identifies a specific developmental process underlying the persistence of these phenotypes. A key strength of the *Drosophila* model is the ability to directly examine airway progenitor niches at defined developmental stages, whereas access to comparable populations in mammalian systems remains technically limited and often restricted to indirect or endpoint analyses. Quantifying progenitor expansion in vivo therefore provides mechanistic insight into developmental origins of airway dysfunction that is ethically difficult to obtain in humans. Together, these findings suggest that disruption of airway progenitor renewal is an early developmental event contributing to long-term impairment of airway function, consistent with the concept that early-life disturbances shape adult health outcomes (*8, 16, 61*). At the molecular level, combined transcriptomic analyses of tracheal tissue and stem-cell niches reveal a previously unrecognized convergence between developmental and immune regulatory programs. In particular, consistent downregulation of Relish-dependent antimicrobial and immune response genes across datasets suggests that innate immune signalling not only mediates defence but also participates in coordinating epithelial growth and progenitor dynamics.

Sex-specific responses to prenatal smoke exposure have been reported in both human and experimental studies, with female offspring often showing greater respiratory or developmental impairment (*62–65*). In females of our study, enrichment of stress-related pathways, including heat shock proteins, may indicate increased cellular vulnerability (*66, 67*). In contrast, males show induction of detoxification-associated genes, such as cytochrome P450 pathways, consistent with adaptive responses to xenobiotic stress (*68, 69*). Together, these patterns suggest that while both sexes experience reduced physiological capacity, females may mount a more constrained stress response, whereas males exhibit broader transcriptional adaptation, potentially reflecting differences in developmental plasticity or stress sensitivity during airway development. One possible explanation is that males may be more sensitive to developmental stress, consistent with previous reports of lower stress resistance in male *Drosophila*. This increased sensitivity may require pronounced transcriptional changes in response to epithelial disruption during airway development.

Several human epidemiological studies have reported transgenerational adverse effects of maternal cigarette smoking on the respiratory health of grandchildren (*70, 71*). In mammals, gestational exposures can directly affect fetal germ cells, thereby influencing thereby both the F_1_ and F_2_ generations through intergenerational effects. Accordingly, the F_3_ (“great-grandchildren”) generation represents the first non-exposed generation. In contrast, in *Drosophila*, embryos develop extracorporally, such that the F_2_ generation is already not subject to indirect exposure and therefore reflects true transgenerational effects. In our study, developmental alterations induced by maternal nicotine exposure were not restricted to the directly exposed F_1_ generation but persisted in subsequent descendants, indicating that early-life exposure can have lasting, multigenerational effects. Notably, the severity of the phenotypes examined diminished across generations down the investigated female line, suggesting a progressive restoration of developmental stability rather than stable transgenerational transmission. This observation is important, as it indicates that early regulatory disruption by preconception e-nicotine can influence subsequent generations without establishing a permanent transgenerational state.

Importantly, although paternal exposure to e-nicotine under the same experimental conditions did not reproduce airway defects in offspring, this finding should be interpreted with caution (*72*). In *Drosophila*, spermatogenesis is largely completed within the first few days after eclosion, and sperm are already mature by approximately 4–5 days of age (*73*). Exposure during only one day may therefore have limited capacity to alter the germline state. These considerations suggest that longer exposure of young males covering the full sperm maturation cycle may have greater potential to influence offspring development.

More broadly, a central strength of this study is the direct examination of airway progenitor cell niches during defined stages of development. Access to these progenitor populations enabled analysis of early cellular processes that are challenging to investigate in mammalian systems, where comparable progenitor populations are less accessible and developmental timing is harder to control experimentally.

Several limitations should be considered. Although *Drosophila* provides a powerful platform to investigate developmental and intergenerational mechanisms, its airway system differs structurally and physiologically from the mammalian lung, and therefore direct extrapolation to human respiratory disease requires caution. In addition, while reduced Relish signalling was associated with nicotine-induced airway defects, this pathway is unlikely to act in isolation, and additional systemic regulators may contribute to the observed phenotypes. Finally, although transgenerational effects were detected across multiple generations, the molecular mechanisms responsible for transmitting these effects remain to be defined.

In summary, our findings identify maternal e-nicotine exposure prior to conception as a determinant of airway development and physiological performance in offspring. By linking early progenitor dysfunction to persistent structural and functional abnormalities, this study provides a developmental explanation for how preconception exposures can influence long-term health. The persistence of these phenotypes across generations highlights early developmental stages as a sensitive window for environmental risk and underscores the importance of considering parental exposures before conception. Together, these findings support a model in which maternal preconception nicotine exposure suppresses Relish-dependent innate immune signalling and limits progenitor expansion within the airway stem-cell niche, leading to structural airway vulnerability and reduced physiological resilience across generations. More broadly, these results position the preconception period as a biologically sensitive window capable of shaping developmental trajectories and disease susceptibility later in life.

## Materials and Methods

| Primary antibodies: | Vendor |
| --- | --- |
| α-phospho-Histone H3 (Ser10), rabbit | Cell Signaling Technology (Danvers, USA)<br>Cat # 53348T |
| α-Dcp1, rabbit | Cell Signaling Technology (Danvers, USA)<br>Cat # 9578S |
| α-Coracle, mouse (C615.16) | DSHB (Iowa City, USA) RRID: AB_1161644 |
| Secondary antibodies: |  |
| α-mouse-AlexaFluor 488, goat | Jackson ImmunoResearch Laboratories (West Grove, USA) |
| α-rabbit-AlexaFluor 488, goat | Life Technologies (Carlsbad, USA) |
| α-mouse-AlexaFluor 555, goat | Cell Signaling Technology (Danvers, USA) |
| α-rabbit-Cy3 | Jackson ImmunoResearch Laboratories (West Grove, USA) |
| Fluorescent dyes |  |
| Roti®-Mount FluorCare DAPI (4,6-diamidino-2-phenylindole) | Carl Roth (Karlsruhe, Germany) |
| Chemicals |  |
| Nicotine ≥ 99% (GC) | Sigma-Aldrich Chemie GmbH (Germany) |

| Experimental models: Organisms/strains | Vendor |
| --- | --- |
| *Drosophila melanogaster* Canton S | BDSC (64349) |
| *Drosophila melanogaster* UAS-Rel-RNAi | BDSC (33661) |
| *Drosophila melanogaster* Control line for RNAi | BDSC (36303) |
| *Drosophila melanogaster* Ci-gal4, UAS-GFP; gal80ts | Xavier Franch-Marro, Barcelona, Spain (74) |
| *Drosophila melanogaster* btl-Gal4, UAS-GFP | Maria Leptin Group, Heidelberg, Germany (75) |
| *Drosophila melanogaster* DSRF-Gal4, UAS-GFP | Jordi Casanova, Barcelona, Spain (76) |

### Experimental models: Drosophila

The following transgenic *Drosophila* strains used were supplied by the Bloomington *Drosophila* Stock Center (BDSC): Canton S (BDSC 64349), *UAS-Rel-RNAi* (BDSC 33661). The following stocks used were generously provided by scientists in the *Drosophila* research community: *btl-Gal4, UAS-GFP* (Maria Leptin, Heidelberg, Germany) (*75*), *Ci-gal4, UAS-GFP; gal80ts* (Xavier Franch-Marro, Barcelona, Spain) (*74*), *DSRF-Gal4, UAS-GFP* (Jordi Casanova, Barcelona, Spain) (*76*).

### Fly crosses

For tissue-specific gene expression in the tracheal system the binary Gal4/UAS system was used (*77*). All flies were maintained on standard cornmeal medium at 25°C under a 12-hour light/dark cycle unless otherwise specified. For trachea-specific knockdown experiments, virgin females carrying the driver line *btl-Gal4, UAS-GFP* lines were crossed with males carrying *UAS-Rel-RNAi* or the corresponding RNAi control line. F_1_ larvae were collected for experiments at the third-instar stage. Male and female larvae were separated based on gonadal morphology under a stereomicroscope and analyzed independently. All procedures were performed in compliance with institutional and state animal welfare regulations (Schleswig-Holstein, Germany). For terminal cell specific knockdown experiments, virgin females carrying the driver line *DSRF-Gal4, UAS-GFP* were crossed with males carrying *UAS-Rel-RNAi* or the RNAi control line.

### Maternal preconception e-nicotine exposure in Drosophila

A custom-built e-nicotine exposure apparatus was designed to deliver controlled vaporized nicotine to *Drosophila* adults. The system consisted of a 175-ml glass vial bisected by a nylon mesh insert to prevent direct heat exposure to the flies. The upper opening was sealed with a foam plug, while the bottom was connected to a coiled nichrome wire (3.7 V, 1.9 A) serving as a heating element as described earlier (*8*).

In brief, virgin female flies (100 per group) were randomly assigned to either vaporized nicotine or vaporized water (control). To minimize stress effects, flies were maintained without CO׈ anaesthesia for at least 5 days before exposure. For each exposure, 10 µl of nicotine solution (5 mM in dH׈O) or dH׈O alone was pipetted onto the heated nichrome coil and vaporized for 15 s. The flies were left in the exposure chamber for 2 min to ensure full inhalation of the vapor and then transferred back to their culture vials. This exposure procedure was repeated hourly for eight cycles. Following the final exposure, e-nicotine-treated and control females were placed on separate grapefruit agar (GA) plates and mated with age-matched nonexposed males (100 per group). Eggs were collected 24hrs later to generate the Fu progeny for downstream analyses.

### Paternal exposure to e-nicotine

5-day-old male flies were subjected to the identical e-nicotine exposure protocol as described above for females. Following exposure, males were crossed with 5-day-old virgin females exposed to ambient air, to generate F׈ offspring for phenotypic analyses.

### Transgenerational breeding strategy

To investigate transgenerational effects, F׈ offspring derived from maternal e-nicotine exposure were grown until age 5 days. Virgin females were collected and then crossed with untreated 5-day-old males from untreated F_0_ to generate the F׈ generation. Control F׈ progeny were obtained by crossing age-matched F׈ virgin females with males whose mothers had been exposed to water. For F׈ generation experiments, the same breeding strategy was applied, using F׈ females derived from each group crossed with males from untreated F_0_. To further assess persistence of early developmental phenotypes, Fu embryos were collected from Fu females using the same crossing scheme.

### Morphometric analysis of embryos, larvae, and adult flies

Egg, larval, and adult body dimensions were quantified using standardized morphometric procedures.

Egg length was measured using synchronized embryos collected over an 8 h laying period. Eggs were transferred onto glass slides and imaged under bright-field microscopy. Egg length was defined as the distance from the anterior to the posterior pole.

For larval measurements, third-instar larvae were collected and heat-killed at 70 °C for 10 s prior to imaging. Larvae were mounted in glycerol on glass slides, aligned along the anterior–posterior axis, and imaged using a stereomicroscope. Body length was measured from the anterior to the posterior end.

Adult body length was measured in flies anesthetized with CO׈ and immobilized on ice during imaging. Flies were positioned laterally, and body length was defined as the distance from the head to the tip of the abdomen.

All morphometric measurements were performed using ImageJ software.

### Assessment of developmental progression

Developmental progression was assessed by monitoring the transition from embryonic to pupal and adult stages under standard culture conditions. The number of pupae was recorded on day 9, and the number of eclosed adult flies was recorded on day 15. Developmental success was expressed as the proportion of eggs reaching pupal and adult stages, respectively, consistent with established developmental timing assays.

### Measurement of tracheal epithelial thickness

On day 5 of larval development, isolated tracheae were mounted in Ibidi mounting medium on glass slides, covered with a coverslip, and sealed with clear nail polish. Images were acquired using an Olympus BX51 microscope equipped with a DP25 camera (Olympus GmbH, Hamburg, Germany) at 20× magnification. Epithelial thickness was measured at three representative locations along the A6/A7 dorsal trunk and major branches using the Polyline tool in the imaging software, and mean values were used for statistical analysis.

### Immunostaining and imaging of larval tracheae and tracheal stem cell niches

Immunostaining of larval tracheae was performed as previously described (Fehon et al., 1994) (*78*) with minor modifications. Briefly, late third-instar larvae were dissected in PBS to isolate the tracheal system, fixed in 4% paraformaldehyde for 30–45 minutes at room temperature, and washed in PBS. Samples were permeabilized in PBS containing 0.1% Triton X-100 and blocked in PBS supplemented with 10% normal goat serum. Tissues were incubated overnight at 4°C with primary antibodies diluted in blocking buffer, followed by appropriate Alexa Fluor–conjugated secondary antibodies and DAPI counterstaining. Primary antibodies used in this study included mouse anti-Coracle (clone C615.16; Developmental Studies Hybridoma Bank, Iowa City, USA; RRID: AB_1161644), rabbit anti–cleaved *Drosophila* Dcp1 (Cell Signaling Technology; RRID: AB_2721060), and rabbit anti–phospho-histone H3 (Ser10) (Cell Signaling Technology; RRID: AB_331535).

Samples were mounted in ProLong Diamond Antifade (Thermo Fisher Scientific) and imaged using an Olympus SZX16 stereo fluorescence microscope or a Leica fluorescence microscope with z-stack acquisition.

### Measurement of terminal cell morphology

Tracheal terminal cell length and branch number were quantified using ImageJ (NIH) with the NeuronJ plugin (*79*). Each individual branch was traced manually to obtain total branch length and number of arborizations per cell. All measurements were performed under identical settings and analyzed consistently across samples.

### Measurement of the area of tracheoblasts

The tracheae of wandering larvae were dissected and the tracheoblasts were selected for microscopy and quantification using Z-stack analysis. Z-stack images were documented in DIC and GFP channels with the Axio Imager.Z1 with Apo Tome (Zeiss). Measurements were performed using the AxioVision SE64 Rel. 4.9 Software (Zeiss).

### Larval locomotor activity assays

Larval locomotor activity was assessed as a measure of muscular fitness and behavioral performance, following the protocol described by Nichols *et al.* (2012) with minor modifications (*80*).

### Negative geotaxis (gravitaxis) assay

Baseline locomotor capacity in young adult flies (5 days old) was assessed using a negative geotaxis assay. Flies were transferred into assay tubes and allowed to rest for 1 min. The tubes were gently tapped three times to bring flies to the bottom, and the number of flies crossing a 7 cm mark within 10 s was recorded. Each experiment was repeated three times per group with 1 min intervals between trials.

### Locomotor activity and behavioral assays (Zantiks system)

Locomotor activity of adult flies (2–3 weeks old) was monitored using the Zantiks behavioral tracking system. Individual flies were placed into a 96-well plate, with one fly per well containing 200 μl of standard food. To minimize experimental variability, control and e-nicotine groups were tested simultaneously within the same plate.

For circadian analysis, locomotor activity was continuously recorded over 24 h under standard light dark conditions. For baseline activity assessment, flies were recorded for 30 min at Zeitgeber time 0 (ZT0, lights-on) under red light conditions, which are minimally aversive to *Drosophila*. Immediately following this baseline recording, flies were exposed to a brief green light stimulus (2 s), and locomotor activity was recorded for 1 min to assess behavioral responsiveness. Activity values were normalized within each experiment prior to pooling. Each experiment was performed in three independent biological replicates.

### Behavioral data processing

Locomotor activity data were extracted from the Zantiks system and normalized within each experiment prior to statistical analysis. For circadian analysis, activity was quantified over time and compared between groups. For the green light stimulation assay, locomotor activity during the 1-min period following stimulation was quantified as relative locomotor activity. A response index was calculated to assess stimulus-induced behavioral changes relative to prestimulus baseline activity measured within the same individuals.

### Hypoxia tolerance assay

Hypoxia tolerance was assessed in early third-instar larvae. Larvae were separated by sex, and groups of 10 individuals were transferred to food-containing vials and allowed to burrow into the medium. The vials were then placed in a sealed glass chamber connected to an oxygen sensor. Nitrogen gas was introduced to gradually reduce the oxygen concentration to approximately 2%, which was maintained below 3% throughout the exposure period. Larvae were kept under hypoxic conditions for 3 hours and were subsequently returned to normoxic conditions for recovery. Recovery was defined as the resumption of crawling behavior. Larvae that failed to recover locomotion within 2 hours after reoxygenation were scored as dead. Each experiment was performed with at least three independent biological replicates.

### Survival assay

Survival was assessed in both maternal (F_0_) and offspring (F׈) flies following maternal e-nicotine exposure. For offspring survival, newly eclosed F׈ adults were collected within 24 hours after eclosion, separated by sex, and maintained in vials containing standard cornmeal medium (70–110 individuals per vial). Flies were transferred to fresh medium three times per week, and survival was monitored until all individuals had died.

For parental survival, F׈ flies were exposed to vaporized nicotine as described above and maintained under the same conditions.

Deaths were recorded at each transfer, and survival data were analyzed using GraphPad Prism. Flies removed for reasons unrelated to natural death were treated as censored observations.

### Gut integrity

Intestinal barrier integrity was assessed using the dye permeability (“Smurf”) assay. Adult flies were fed food supplemented with a non-absorbable blue dye (FD&C Blue No. 1, 2.5% w/v in 5% sucrose) for 24 hours at 25 °C. Flies were then examined under a stereomicroscope and classified based on dye distribution. Flies in which the dye remained confined to the gut were scored as non-Smurf, whereas flies showing dye leakage beyond the digestive tract were scored as Smurf-positive, indicating compromised barrier integrity. For each experimental group, 20–30 flies per sex were analyzed per replicate, with at least three independent biological replicates. The proportion of Smurf-positive flies was quantified and compared between groups.

### RNA-Seq and data processing

For larval tracheae, tracheal tissues were dissected from early third-instar larvae in ice-cold phosphate-buffered saline under a stereomicroscope. Approximately 40 to 60 tracheae were collected per biological replicate. Total RNA was isolated using the NucleoSpin RNA Kit (Macherey-Nagel) according to the manufacturer’s instructions. RNA quality was assessed using an Agilent Bioanalyzer, and only samples with an RNA integrity number greater than 7 were used for downstream analysis.

For tracheal stem cell niche samples, Tr4/Tr5 niches were dissected from late third-instar larvae of *Ci-Gal4, UAS-GFP; Gal80*^ts^ flies under fluorescence guidance. Owing to the limited RNA input obtained from isolated niches, cDNA synthesis and amplification were performed using a Smart-seq3 protocol before library preparation (*81, 82*).

For both sample types, poly(A)+ RNA was enriched, fragmented, and used for strand-specific cDNA library preparation with random priming. Libraries were sequenced as 150-bp paired-end reads on an Illumina NextSeq 500 platform by Eurofins Genomics. RNA-Seq data were processed on the European Galaxy Server (https://usegalaxy.eu). The workflow included read quality assessment, adapter and quality trimming, alignment, and gene-level counting (*83*). Reads were aligned to the *Drosophila* reference genome using STAR (*84*), and gene counts were generated with featureCounts (*85*) using the corresponding genome annotation. For read counting, a minimum mapping quality of 12 and a minimum overlap of 1 bp with an annotated exon were required. Differential expression analysis was performed from gene count tables using DESeq2 (*86*), and adjusted p values (padj) were used to define differentially expressed genes.

### Gene set enrichment analysis

For functional enrichment, we performed gene set enrichment analysis (GSEA) in R using Gene Ontology Biological Process (GO BP) gene sets. Genes were ranked based on the RNA Seq differential expression results, and enrichment was tested using a preranked GSEA framework. Significantly enriched GO BP terms were identified based on multiple testing adjusted p values, and the top 10 most significant GO BP terms were reported separately for female and male comparisons.

### Protein-protein interaction (PPI) network analysis

Protein-protein interactions among the DEGs were retrieved from the STRING database (version 12.0; accessed via the web interface) using *Drosophila* as the reference organism and applying a high-confidence interaction score threshold (≥ 0.7). Both experimentally validated and database-curated interactions were included. The resulting interaction table was imported into Cytoscape (version 3.10.3) for network visualization.

## Supporting information

Supplementary file_Preconception e-nicotine

## Data availability

The sequencing data are available at EGA under accession XXX (submission ongoing).

## Quantitative and statistical analysis

All experiments were performed with at least three independent biological replicates. All statistical analyses were performed using GraphPad Prism (version 10). Data are presented as mean ± SEM unless otherwise indicated. Sample sizes are specified in the figure legends and refer to the relevant experimental unit for each assay. Comparisons between two groups were performed using unpaired two-tailed Student’s t-tests when data were normally distributed. For comparisons involving multiple groups, one-way or two-way analysis of variance (ANOVA) followed by appropriate post hoc tests was applied, as indicated in the figure legends. Survival data were analyzed using the Kaplan–Meier method and compared using the log-rank test. Differences were considered statistically significant at *P* < 0.05. For transcriptomic analyses, differential gene expression was determined using an adjusted P-value threshold (padj < 0.05) following multiple-testing correction. Images were acquired using Olympus cellSens and Leica Application Suite software. Image analysis was performed using ImageJ and NeuronJ plugins.

## Acknowledgements

We thank Britta Laubenstein and Christiane Sandberg for excellent technical assistance. We also thank the DZL Disease Spanning Working Group “Lung-Environment Interaction” for fruitful discussion of the project.

## Funding

The present work was supported by the award of the Balzan Prize to the German Center for Lung Research (DZL) scientists Erika von Mutius, Klaus F. Rabe, Werner Seeger and Tobias Welte and further by the Leibniz Science Campus Evolutionary Medicine of the Lung (EvoLUNG). Qiaoyan Tan was supported by the China Scholarship Council (Grant number 202508500095). Göksel DOĞAN was supported by the TUBITAK-2219 fellowship program (2219-International Postdoctoral Research Fellowship Program for Turkish Citizens).

## Notes

### Competing Interest Statement

The authors have declared no competing interest.

## References

1. Z. Zhao et al., E-cigarette use among adults in China: findings from repeated cross-sectional surveys in 2015-16 and 2018-19. Lancet Public Health 5, e639–e649 (2020).

2. J. M. Kinnunen et al., Electronic cigarette use among 14- to 17-year-olds in Europe. Eur J Public Health 31, 402–408 (2021).

3. P. D. Gluckman, M. A. Hanson, T. Buklijas, A conceptual framework for the developmental origins of health and disease. J Dev Orig Health Dis 1, 6–18 (2010).

4. J. J. Heindel et al., Developmental Origins of Health and Disease: Integrating Environmental Influences. Endocrinology 156, 3416–3421 (2015).

5. T. P. Fleming et al., Origins of lifetime health around the time of conception: causes and consequences. Lancet 391, 1842–1852 (2018).

6. B. Liu et al., National Estimates of e-Cigarette Use Among Pregnant and Nonpregnant Women of Reproductive Age in the United States, 2014-2017. JAMA Pediatr 173, 600–602 (2019).

7. J. F. Bertholon, M. H. Becquemin, I. Annesi-Maesano, B. Dautzenberg, Electronic cigarettes: a short review. Respiration 86, 433–438 (2013).

8. N. El-Merhie et al., Sex dependent effect of maternal e-nicotine on F1 Drosophila development and airways. Sci Rep 11, 4441 (2021).

9. W. Luck, H. Nau, R. Hansen, R. Steldinger, Extent of nicotine and cotinine transfer to the human fetus, placenta and amniotic fluid of smoking mothers. Dev Pharmacol Ther 8, 384–395 (1985).

10. X. Wang, N. L. Lee, I. Burstyn, Smoking and use of electronic cigarettes (vaping) in relation to preterm birth and small-for-gestational-age in a 2016 U.S. national sample. Prev Med 134, 106041 (2020).

11. V. M. Cardenas, M. M. Ali, L. A. Fischbach, W. N. Nembhard, Dual use of cigarettes and electronic nicotine delivery systems during pregnancy and the risk of small for gestational age neonates. Ann Epidemiol 52, 86–92 e82 (2020).

12. Y. Zhang et al., Maternal electronic cigarette exposure in relation to offspring development: a comprehensive review. Am J Obstet Gynecol MFM 4, 100659 (2022).

13. E. K. Nanninga et al., Adverse Maternal and Infant Outcomes of Women Who Differ in Smoking Status: E-Cigarette and Tobacco Cigarette Users. Int J Environ Res Public Health 20, (2023).

14. L. G. Rollins et al., Electronic Cigarette Use During Preconception and/or Pregnancy: Prevalence, Characteristics, and Concurrent Mental Health Conditions. J Womens Health (Larchmt) 29, 780–788 (2020).

15. C. Svanes, J. W. Holloway, S. Krauss-Etschmann, Preconception origins of asthma, allergies and lung function: The influence of previous generations on the respiratory health of our children. J Intern Med 293, 531–549 (2023).

16. C. Wongtrakool, N. Wang, D. M. Hyde, J. Roman, E. R. Spindel, Prenatal nicotine exposure alters lung function and airway geometry through alpha7 nicotinic receptors. Am J Respir Cell Mol Biol 46, 695–702 (2012).

17. K. L. Sandberg, K. E. Pinkerton, S. D. Poole, P. A. Minton, H. W. Sundell, Fetal nicotine exposure increases airway responsiveness and alters airway wall composition in young lambs. Respir Physiol Neurobiol 176, 57–67 (2011).

18. M. J. Blacquiere et al., Maternal smoking during pregnancy induces airway remodelling in mice offspring. Eur Respir J 33, 1133–1140 (2009).

19. H. S. Sekhon et al., Prenatal nicotine increases pulmonary alpha7 nicotinic receptor expression and alters fetal lung development in monkeys. J Clin Invest 103, 637–647 (1999).

20. E. R. Spindel, C. T. McEvoy, The Role of Nicotine in the Effects of Maternal Smoking during Pregnancy on Lung Development and Childhood Respiratory Disease. Implications for Dangers of E-Cigarettes. Am J Respir Crit Care Med 193, 486–494 (2016).

21. A. Gambadauro et al., Impact of E-Cigarettes on Fetal and Neonatal Lung Development: The Influence of Oxidative Stress and Inflammation. Antioxidants (Basel) 14, (2025).

22. A. P. Tackett et al., Prospective study of e-cigarette use and respiratory symptoms in adolescents and young adults. Thorax 79, 163–168 (2024).

23. A. Alnajem et al., Use of electronic cigarettes and secondhand exposure to their aerosols are associated with asthma symptoms among adolescents: a cross-sectional study. Respir Res 21, 300 (2020).

24. B. W. Chaffee et al., E-cigarette use and adverse respiratory symptoms among adolescents and Young adults in the United States. Prev Med 153, 106766 (2021).

25. A. Noel et al., In utero exposures to electronic-cigarette aerosols impair the Wnt signaling during mouse lung development. Am J Physiol Lung Cell Mol Physiol 318, L705–L722 (2020).

26. A. Noel et al., Sex-Specific Alterations of the Lung Transcriptome at Birth in Mouse Offspring Prenatally Exposed to Vanilla-Flavored E-Cigarette Aerosols and Enhanced Susceptibility to Asthma. Int J Environ Res Public Health 20, (2023).

27. D. M. Aslaner et al., E-cigarette vapor exposure in utero causes long-term pulmonary effects in offspring. Am J Physiol Lung Cell Mol Physiol 323, L676–L682 (2022).

28. K. M. Cahill et al., In utero exposures to mint-flavored JUUL aerosol impair lung development and aggravate house dust mite-induced asthma in adult offspring mice. Toxicology 477, 153272 (2022).

29. M. R. Orzabal et al., Impact of E-cig aerosol vaping on fetal and neonatal respiratory development and function. Transl Res 246, 102–114 (2022).

30. M. Affolter, E. Caussinus, Tracheal branching morphogenesis in Drosophila: new insights into cell behaviour and organ architecture. Development 135, 2055–2064 (2008).

31. A. Ghabrial, S. Luschnig, M. M. Metzstein, M. A. Krasnow, Branching morphogenesis of the Drosophila tracheal system. Annu Rev Cell Dev Biol 19, 623–647 (2003).

32. F. Chen, M. A. Krasnow, Progenitor outgrowth from the niche in Drosophila trachea is guided by FGF from decaying branches. Science 343, 186–189 (2014).

33. K. T. Sirocko et al., Early-life exposure to tobacco smoke alters airway signaling pathways and later mortality in D. melanogaster. Environ Pollut 309, 119696 (2022).

34. L. Blackie et al., The sex of organ geometry. Nature 630, 392–400 (2024).

35. H. H. Kampinga, E. A. Craig, The HSP70 chaperone machinery: J proteins as drivers of functional specificity. Nat Rev Mol Cell Biol 11, 579–592 (2010).

36. C. Samakovlis, D. A. Kimbrell, P. Kylsten, A. Engstrom, D. Hultmark, The immune response in Drosophila: pattern of cecropin expression and biological activity. EMBO J 9, 2969–2976 (1990).

37. C. Persson, S. Oldenvi, H. Steiner, Peptidoglycan recognition protein LF: a negative regulator of Drosophila immunity. Insect Biochem Mol Biol 37, 1309–1316 (2007).

38. N. Basbous et al., The Drosophila peptidoglycan-recognition protein LF interacts with peptidoglycan-recognition protein LC to downregulate the Imd pathway. EMBO Rep 12, 327–333 (2011).

39. N. Arquier, M. Bourouis, J. Colombani, P. Leopold, Drosophila Lk6 kinase controls phosphorylation of eukaryotic translation initiation factor 4E and promotes normal growth and development. Curr Biol 15, 19–23 (2005).

40. B. Lemaitre, J. M. Reichhart, J. A. Hoffmann, Drosophila host defense: differential induction of antimicrobial peptide genes after infection by various classes of microorganisms. Proc Natl Acad Sci U S A 94, 14614–14619 (1997).

41. A. Kleino et al., Pirk is a negative regulator of the Drosophila Imd pathway. J Immunol 180, 5413–5422 (2008).

42. J. M. Park et al., Targeting of TAK1 by the NF-kappa B protein Relish regulates the JNK-mediated immune response in Drosophila. Genes Dev 18, 584–594 (2004).

43. R. S. Khush, F. Leulier, B. Lemaitre, Drosophila immunity: two paths to NF-kappaB. Trends Immunol 22, 260–264 (2001).

44. M. S. Busse, C. P. Arnold, P. Towb, J. Katrivesis, S. A. Wasserman, A kappaB sequence code for pathway-specific innate immune responses. EMBO J 26, 3826–3835 (2007).

45. M. Weaver, M. A. Krasnow, Dual origin of tissue-specific progenitor cells in Drosophila tracheal remodeling. Science 321, 1496–1499 (2008).

46. C. de Miguel, J. Cruz, D. Martin, X. Franch-Marro, Dual role of FGF in proliferation and endoreplication of Drosophila tracheal adult progenitor cells. J Mol Cell Biol 12, 32–41 (2020).

47. M. J. Hendzel et al., Mitosis-specific phosphorylation of histone H3 initiates primarily within pericentromeric heterochromatin during G2 and spreads in an ordered fashion coincident with mitotic chromosome condensation. Chromosoma 106, 348–360 (1997).

48. Z. Song, K. McCall, H. Steller, DCP-1, a Drosophila cell death protease essential for development. Science 275, 536–540 (1997).

49. S. T. Holgate, The airway epithelium is central to the pathogenesis of asthma. Allergol Int 57, 1–10 (2008).

50. D. J. Tschumperlin, J. D. Shively, T. Kikuchi, J. M. Drazen, Mechanical stress triggers selective release of fibrotic mediators from bronchial epithelium. Am J Respir Cell Mol Biol 28, 142–149 (2003).

51. H. S. Sekhon, J. A. Keller, N. L. Benowitz, E. R. Spindel, Prenatal nicotine exposure alters pulmonary function in newborn rhesus monkeys. Am J Respir Crit Care Med 164, 989–994 (2001).

52. C. Mwase et al., Mechanical Compression of Human Airway Epithelial Cells Induces Release of Extracellular Vesicles Containing Tenascin C. Cells 11, (2022).

53. F. D. Gilliland, Y. F. Li, J. M. Peters, Effects of maternal smoking during pregnancy and environmental tobacco smoke on asthma and wheezing in children. Am J Respir Crit Care Med 163, 429–436 (2001).

54. G. S. Maritz, R. Harding, Life-long programming implications of exposure to tobacco smoking and nicotine before and soon after birth: evidence for altered lung development. Int J Environ Res Public Health 8, 875–898 (2011).

55. J. C. Hendricks et al., Rest in Drosophila is a sleep-like state. Neuron 25, 129–138 (2000).

56. T. A. Slotkin et al., Developmental Neurotoxicity of Tobacco Smoke Directed Toward Cholinergic and Serotonergic Systems: More Than Just Nicotine. Toxicol Sci 147, 178–189 (2015).

57. S. Godleski, S. Shisler, K. Colton, M. Leising, Prenatal Tobacco Exposure and Behavioral Disorders in Children and Adolescents: Systematic Review and Meta-Analysis. Pediatr Rep 16, 736–752 (2024).

58. C. Hetru, J. A. Hoffmann, NF-kappaB in the immune response of Drosophila. Cold Spring Harb Perspect Biol 1, a000232 (2009).

59. A. Di Stefano et al., Increased expression of nuclear factor-kappaB in bronchial biopsies from smokers and patients with COPD. Eur Respir J 20, 556–563 (2002).

60. C. Y. Tsai, H. C. Chou, C. M. Chen, Perinatal nicotine exposure alters lung development and induces HMGB1-RAGE expression in neonatal mice. Birth Defects Res 113, 570–578 (2021).

61. P. D. Gluckman, M. A. Hanson, C. Cooper, K. L. Thornburg, Effect of in utero and early-life conditions on adult health and disease. N Engl J Med 359, 61–73 (2008).

62. R. Alati, A. Al Mamun, M. O’Callaghan, J. M. Najman, G. M. Williams, In utero and postnatal maternal smoking and asthma in adolescence. Epidemiology 17, 138–144 (2006).

63. K. D. Reddy, B. G. G. Oliver, Sex-specific effects of in utero and adult tobacco smoke exposure. Am J Physiol Lung Cell Mol Physiol 320, L63–L72 (2021).

64. W. Wei et al., Association of maternal smoking during pregnancy with youth depression and subsequent adult chronic diseases in offspring. Transl Psychiatry 16, (2026).

65. B. Wang et al., Offspring sex affects the susceptibility to maternal smoking-induced lung inflammation and the effect of maternal antioxidant supplementation in mice. J Inflamm (Lond) 17, 24 (2020).

66. M. E. Feder, G. E. Hofmann, Heat-shock proteins, molecular chaperones, and the stress response: evolutionary and ecological physiology. Annu Rev Physiol 61, 243–282 (1999).

67. J. Tower, Heat shock proteins and Drosophila aging. Exp Gerontol 46, 355–362 (2011).

68. G. Le Goff, F. Hilliou, Resistance evolution in Drosophila: the case of CYP6G1. Pest Manag Sci 73, 493–499 (2017).

69. S. F. Zhou, J. P. Liu, B. Chowbay, Polymorphism of human cytochrome P450 enzymes and its clinical impact. Drug Metab Rev 41, 89–295 (2009).

70. G. M. Mahon, G. H. Koppelman, J. M. Vonk, Grandmaternal smoking, asthma and lung function in the offspring: the Lifelines cohort study. Thorax 76, 441–447 (2021).

71. Y. F. Li, B. Langholz, M. T. Salam, F. D. Gilliland, Maternal and grandmaternal smoking patterns are associated with early childhood asthma. Chest 127, 1232–1241 (2005).

72. C. Svanes et al., Father’s environment before conception and asthma risk in his children: a multi-generation analysis of the Respiratory Health In Northern Europe study. Int J Epidemiol 46, 235–245 (2017).

73. L. Fabian, J. A. Brill, Drosophila spermiogenesis: Big things come from little packages. Spermatogenesis 2, 197–212 (2012).

74. K. Campbell, G. Lebreton, X. Franch-Marro, J. Casanova, Differential roles of the Drosophila EMT-inducing transcription factors Snail and Serpent in driving primary tumour growth. PLoS Genet 14, e1007167 (2018).

75. J. Bossen et al., Adult and Larval Tracheal Systems Exhibit Different Molecular Architectures in Drosophila. Int J Mol Sci 24, (2023).

76. L. Gervais, J. Casanova, The Drosophila homologue of SRF acts as a boosting mechanism to sustain FGF-induced terminal branching in the tracheal system. Development 138, 1269–1274 (2011).

77. A. H. Brand, N. Perrimon, Targeted gene expression as a means of altering cell fates and generating dominant phenotypes. Development 118, 401–415 (1993).

78. R. G. Fehon, I. A. Dawson, S. Artavanis-Tsakonas, A Drosophila homologue of membrane-skeleton protein 4.1 is associated with septate junctions and is encoded by the coracle gene. Development 120, 545–557 (1994).

79. E. Meijering et al., Design and validation of a tool for neurite tracing and analysis in fluorescence microscopy images. Cytometry A 58, 167–176 (2004).

80. C. D. Nichols, J. Becnel, U. B. Pandey, Methods to assay Drosophila behavior. J Vis Exp, (2012).

81. M. Hagemann-Jensen et al., Single-cell RNA counting at allele and isoform resolution using Smart-seq3. Nat Biotechnol 38, 708–714 (2020).

82. H. Meyer et al., Combined transcriptome and proteome profiling reveal cell-type-specific functions of Drosophila garland and pericardial nephrocytes. Commun Biol 7, 1424 (2024).

83. E. Afgan et al., The Galaxy platform for accessible, reproducible and collaborative biomedical analyses: 2018 update. Nucleic Acids Res 46, W537–W544 (2018).

84. A. Dobin et al., STAR: ultrafast universal RNA-seq aligner. Bioinformatics 29, 15–21 (2013).

85. Y. Liao, G. K. Smyth, W. Shi, featureCounts: an efficient general purpose program for assigning sequence reads to genomic features. Bioinformatics 30, 923–930 (2014).

86. M. I. Love, W. Huber, S. Anders, Moderated estimation of fold change and dispersion for RNA-seq data with DESeq2. Genome Biol 15, 550 (2014).

