## Supplementary file_Preconception e-nicotine for "Preconception e-nicotine impairs airway development and progenitor proliferation across generations in Drosophila *Melanogaster*"

**Supplementary Figure 1.** Maternal e-nicotine exposure increases mortality and reduces lifespan in adult females

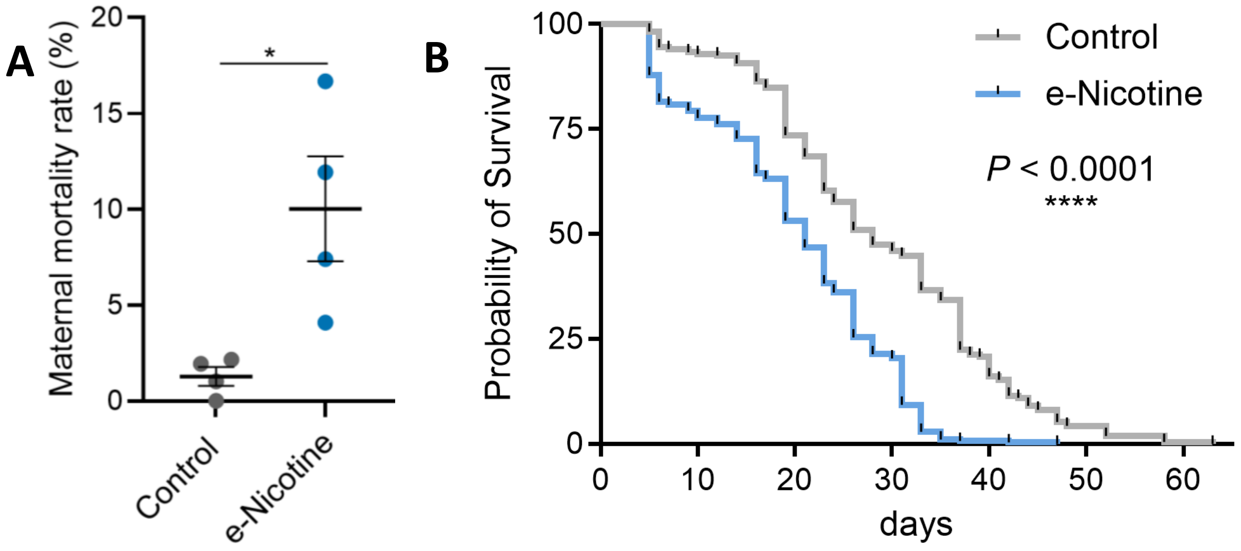

**Supplementary Figure 2.** Paternal e-nicotine exposure affects survival and locomotor performance in $F_0$ flies but does not impair developmental outcomes in $F_1$ offspring

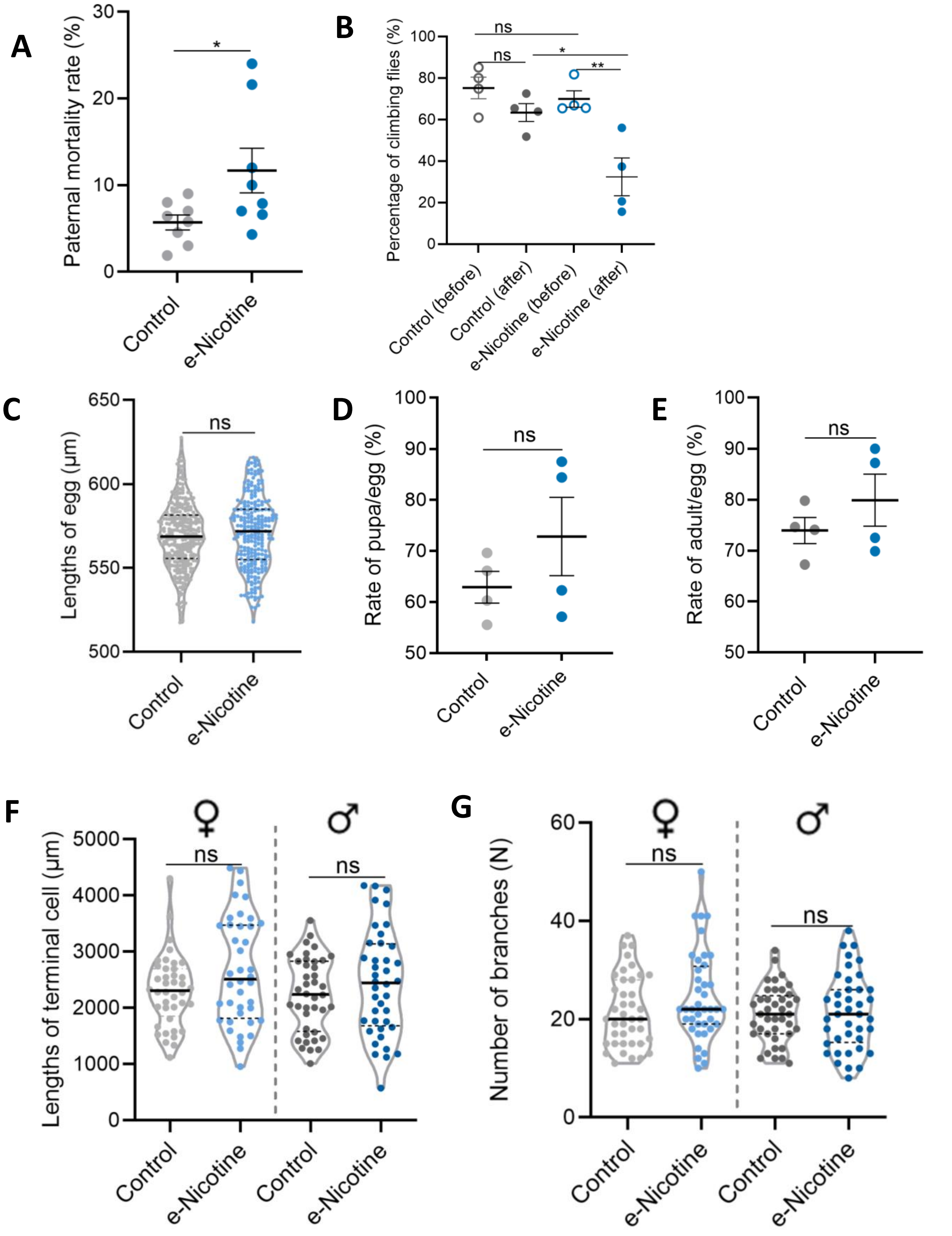

**Supplementary Figure 3**. Maternal e-nicotine exposure alters stimulus-induced behavioral responses in female $F_1$ offspring without affecting baseline locomotor performance

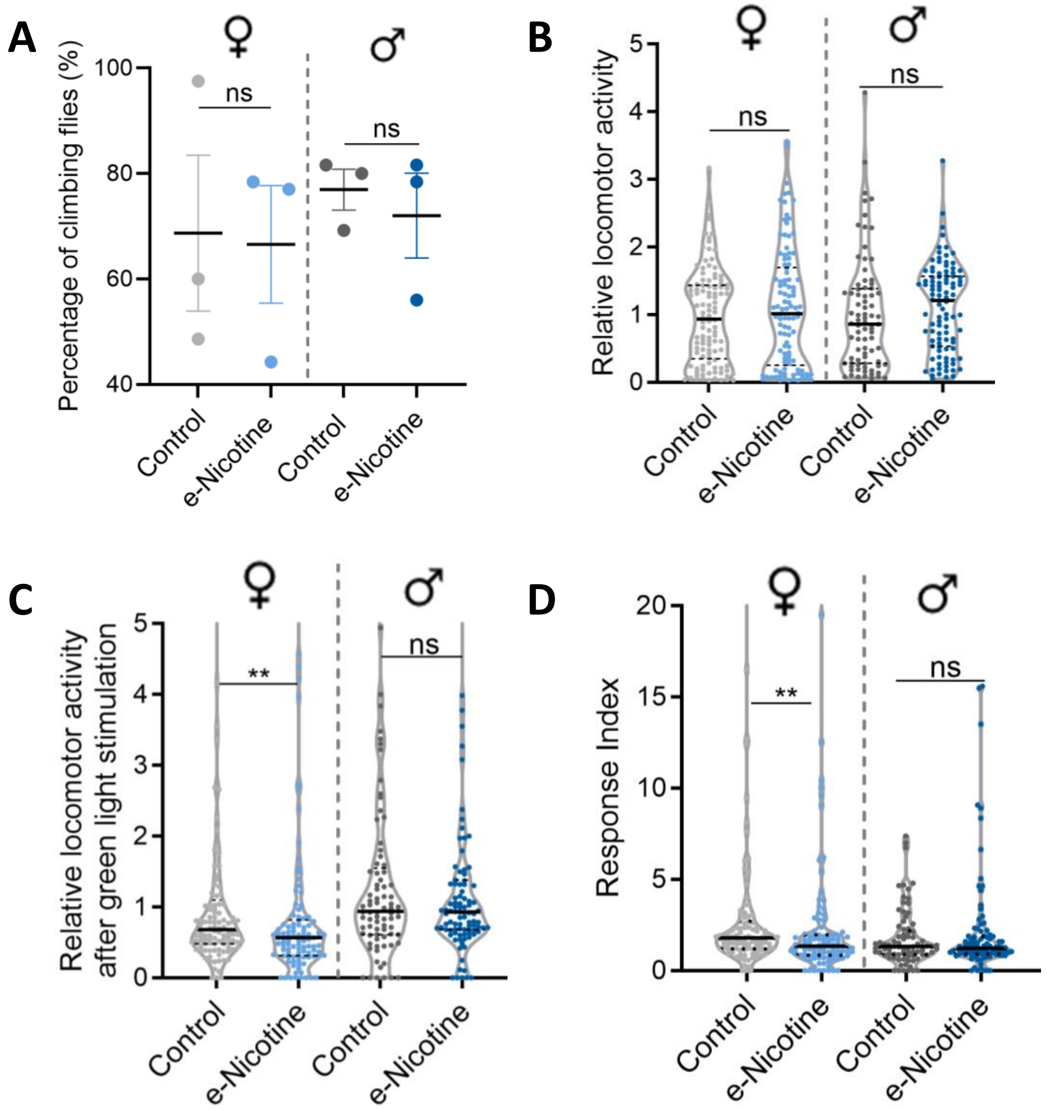

**Supplementary Figure 4.** Preconception e-nicotine exposure reduces Tr4 tracheal progenitor niche area in $F_1$ larvae.

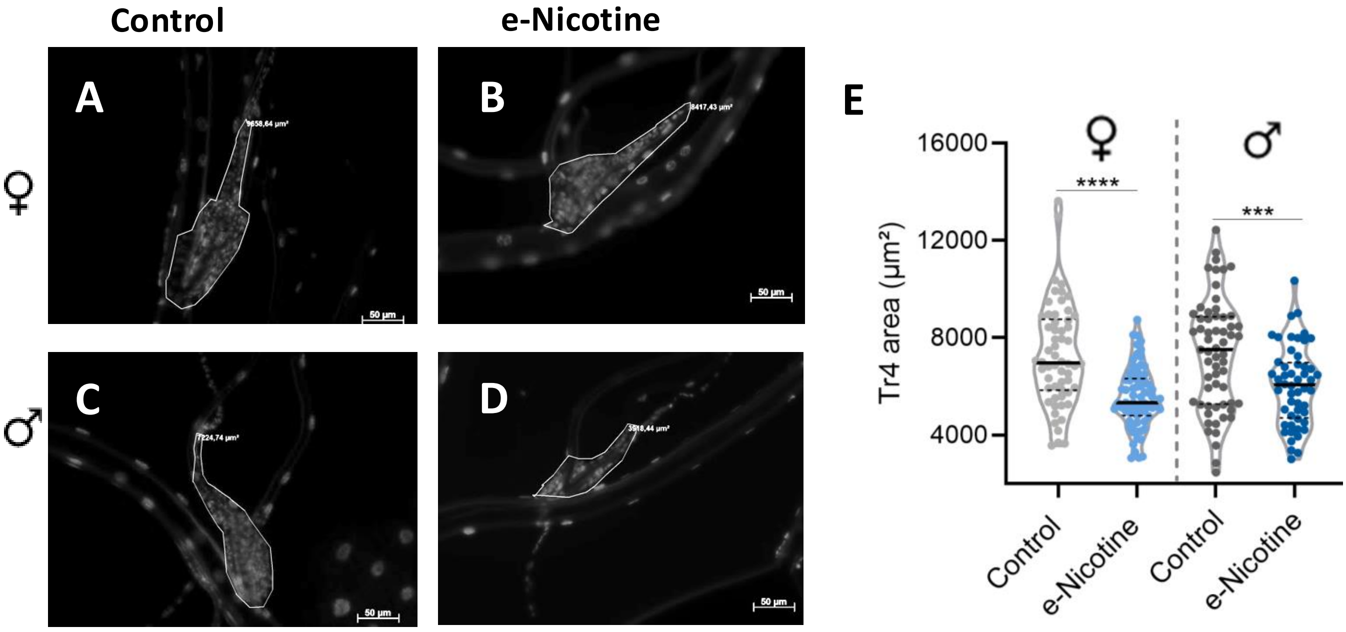

### Supplementary Figure legends:

**Supplementary Figure 1: Maternal e-nicotine exposure increases mortality and reduces lifespan in $F_0$ flies**

(**A**) Mortality rate of $F_0$ adult female flies following e-nicotine exposure. Mortality differed between control and e-nicotine–exposed groups ($P = 0.0473$).

(**B**) Lifespan analysis of $F_0$ female flies exposed to e-nicotine. Median survival was 28 days in controls and 21 days in e-nicotine–exposed flies (n = 294 and 307, respectively; $P < 0.0001$).

Data were obtained from four independent biological experiments for mortality analysis (A) and three independent biological experiments for lifespan analysis (B). Sample sizes were 90–109 flies per group in each experiment. Statistical significance was determined using an unpaired two-tailed Student's t test with Welch's correction for mortality analysis and the log-rank (Mantel–Cox) test for survival analysis. Significance levels are indicated as follows: * ($P < 0.05$), and **** ($P < 0.0001$).

**Supplementary Figure 2: Paternal e-nicotine exposure affects survival and locomotor performance in $F_0$ flies but does not impair developmental or morphological outcomes in $F_1$ offspring.**

(**A**) Mortality rate of $F_0$ adult male flies following paternal e-nicotine exposure. A difference in mortality was observed between control and e-nicotine–exposed groups ($P = 0.0446$).

(**B**) Locomotor activity of F0 adult male flies before and after paternal e-nicotine exposure. Comparisons are shown for e-nicotine flies before versus after exposure ($P = 0.0289$), e-nicotine versus control flies after treatment ($P = 0.0221$), and control flies before versus after treatment ($P = 0.1617$).

(**C**) Egg length in $F_1$ progeny derived from control and paternal e-nicotine–exposed males (569.24 ± 1.11 µm in controls vs. 571.14 ± 1.35 µm in e-nicotine; n = 307 and 247 eggs, respectively; $P = 0.2764$).

(**D** and **E**) Pupariation rate at day 9 (D) and adult emergence rate at day 15 (E) were quantified as indicators of developmental progression and viability ($P = 0.2755$ and $P = 0.3370$, respectively).

(**F** and **G**) Terminal cell length (F) in female offspring ($P = 0.0555$) or male offspring ($P = 0.2544$) and terminal branch number (G) in female offspring ($P = 0.1153$) or male offspring ($P = 0.7660$) derived from both control and paternal e-nicotine–exposed males, n = 40 per group.

Data were obtained from four independent biological experiments unless otherwise indicated (A, eight experiments). Statistical significance was determined using paired or unpaired two-tailed Student's t tests, with Welch's correction applied when variances were unequal. Significance levels are indicated as follows: ns ($P > 0.05$), * ($P < 0.05$), and **($P < 0.01$).

**Supplementary Figure 3: Maternal e-nicotine exposure alters stimulus-induced behavioral responses in female $F_1$ offspring without affecting baseline locomotor performance.**

(**A**) Negative geotaxis performance of female and male $F_1$ offspring derived from control and maternal e-nicotine exposed females. Climbing ability was assessed as the percentage of flies reaching the defined height within the testing period.

(**B**) Baseline locomotor activity of female ($P = 0.5537$) and male $F_1$ offspring ($P = 0.0599$) prior to stimulation. Activity values were quantified under resting conditions.

(**C**) Locomotor activity of female ($P = 0.0062$) and male $F_1$ offspring ($P = 0.9908$), following green light stimulation. Activity values were quantified after stimulation as an indicator of behavioral responsiveness.

(**D**) Response index of female (*P* = 0.0028) and male $F_1$ offspring (*P* = 0.8882), following green light stimulation. The response index was calculated to quantify behavioral responsiveness to stimulation.

Data were obtained from three independent biological experiments. Sample sizes were n = 20 flies per group for geotaxis assays (A) and female control (n = 109), female e-nicotine (n = 102), male control (n = 81), and male e-nicotine (n = 99) for locomotor activity and response analyses (B to D). Statistical significance was determined using the two-tailed Mann–Whitney test. Significance levels are indicated as follows: ns ($P > 0.05$), and **($P < 0.01$).

**Supplementary Figure 4: Preconception e-nicotine exposure reduces Tr4 tracheal progenitor niche area in $F_1$ larvae.**

(**A** to **D**) Representative fluorescence images of the Tr4 tracheal progenitor niche in female (A, B) and male (C, D) $F_1$ larvae derived from control and maternal e-nicotine–exposed groups. Stem cell nuclei were visualized using DAPI staining to define the boundaries of the Tr4 progenitor niche.

(**E**) Quantification of the Tr4 progenitor niche area in female (7281.84 ± 276.43 μm² in controls vs. 5544.40 ± 150.79 μm² in e-nicotine, n = 58 and 70, respectively; $P < 0.0001$) and male $F_1$ larvae (7324.59 ± 292.36 μm² in controls vs. 6032.01 ± 225.96 μm² in e-nicotine, n = 61 and 52, respectively; *P* = 0.0007).

Data were pooled from three independent biological experiments, with more than 15 larvae dissected per group in each experiment. Data are presented as mean ± SEM. Statistical analysis was performed using an unpaired two-tailed t test with Welch's correction.

All scale bars, 50 μm. Significance levels are indicated as follows: *** ($P < 0.001$), and **** ($P < 0.0001$).
